# Trimodal Single-Cell Gene Regulatory Networks Reveal Principles of Stemness Loss and Cell Fate Acquisition in Human Hematopoiesis

**DOI:** 10.1101/2025.09.11.675740

**Authors:** Carmen G. Palii, Steven Tur, Sirui Yan, William J.R. Longabaugh, Romeo Solano, F. Jeffrey Dilworth, Jeffrey A. Ranish, Marjorie Brand

## Abstract

Hematopoiesis requires the coordinated loss of stemness and acquisition of lineage identity, yet the regulatory logic and principles underlying these transitions has remained elusive. Single-cell studies suggest that hematopoietic stem and progenitor cells progress along continuous trajectories, but this view conflicts with the existence of discrete, functionally validated populations. Here, we establish the first dynamic, enhancer-based gene regulatory networks (eGRNs) that resolve the molecular programs underlying early human hematopoietic fate decisions. Built on a high-resolution trimodal framework generated with TEA-seq, these networks integrate simultaneously measured transcription factor abundance, enhancer and promoter accessibility, and gene expression within single-cell trajectories anchored to immunophenotypically defined populations. Our framework reveals that stemness loss and lineage acquisition are temporally and mechanistically uncoupled: stemness programs decline gradually through reduced TF abundance long before chromatin closure, whereas lineage identity emerges stepwise through enhancer reconfiguration and activation of lineage-defining eGRNs. This process generates discrete regulatory states that align with immunophenotypically defined populations. Together, these findings reconcile continuous and discrete models of hematopoiesis and establish eGRNs as a powerful framework for defining cell types by their regulatory logic. In addition, we provide an interactive web-based resource to facilitate further investigation of eGRNs and trajectories during early human hematopoiesis.

## INTRODUCTION

Hematopoiesis is a hierarchical process in which multipotent hematopoietic stem cells (HSCs) give rise to all blood cell types. This dynamic system sustains lifelong blood production while rapidly adapting to environmental changes ^1,2^. As such, hematopoiesis provides an ideal model for dissecting the mechanisms underlying cell fate decisions.

Over the past decade, single-cell technologies profiling the transcriptome (scRNA-seq) ^3^, proteome ^4–6^, or chromatin accessibility (scATAC-seq) ^7^ have transformed our understanding of hematopoiesis. These approaches revealed that differentiation proceeds as a continuum along lineage trajectories rather than through discrete steps, shifting the paradigm from a stepwise to a gradual model of fate commitment ^8–10^. Yet, this high-resolution view has blurred classical immunophenotypic boundaries, raising fundamental questions: What defines a cell type? And which characteristic best captures cellular identity?

Because each single-cell approach (or modality) reflects only one facet of cell state, an integrated view is essential ^11^. However, most multimodal analyses rely on correlating datasets that were measured separately ^12–14^, even though mRNA and protein levels are only weakly correlated ^6,15–19^ and chromatin accessibility often precedes transcription ^20–25^. These discrepancies limit interpretability and highlight the need to measure multiple modalities simultaneously in the same single cells. While dual-modality approaches like CITE-seq (measures RNA and cell surface proteins) ^26,27^ and 10x Multiome/SNARE-seq (measures RNA and chromatin accessibility) ^28^ are now widely used, only TEA-seq ^29^ and DOGMA-seq ^30^ capture RNA, chromatin and surface proteins in the same single cells.

Beyond mapping trajectories, the central challenge is to uncover the molecular logic and principles of cell fate transitions. Fluctuations in the expression of lineage-specifying transcription factors (LS-TFs) can influence fate ^4,10,31,32^, yet lineage-tracing with scRNA-seq shows that fate choices arise earlier than transcriptomes alone can predict ^33^. This points to chromatin accessibility ^34^ and enhancer-mediated regulation ^35^ as critical upstream determinants of fate. Because LS-TFs are often redeployed across lineages ^1,36^, cell identity is not dictated by TFs or enhancers in isolation, but by their combinations within gene regulatory networks (GRNs) ^37–41^. Although GRNs of hematopoiesis have been described ^19,22,42–44^, none have integrated gene expression with enhancer accessibility along immunophenotypically aware trajectories. As a result, the regulatory logic of early hematopoietic cell fate decisions has remained unresolved.

Here, by applying probabilistic modeling to trimodal TEA-seq profiles of RNA, chromatin, and surface proteins measured in the same single cells along a differentiation time course, we construct time-ordered differentiation paths and enhancer-based GRNs at unprecedented resolution. This framework refines our understanding of cell identity, uncovers how transcription factor fluctuations and chromatin remodeling act as temporally distinct drivers of fate, and provides the community with a web-based, interactive resource to explore enhancer-based GRNs and human hematopoietic trajectories.

## RESULTS

### TEAseq Analysis of Early Human Hematopoietic Differentiation

Hematopoiesis has been extensively studied using single-cell assays that profile distinct modalities, such as the transcriptome (scRNA-seq) ^3^, chromatin accessibility (scATAC-seq) ^7^, or cell surface proteins (CyTOF) ^4,5^. However, these modalities are typically measured independently, each providing only a partial view of the regulatory landscape.

To overcome this limitation, we used a trimodal approach (TEAseq) for single-cell analysis of mRNA **<u>t</u>**ranscripts, surface protein **<u>e</u>**pitopes and chromatin **<u>a</u>**ccessibility in which all three modalities are measured simultaneously in the same single cells^29^. Primary human CD34⁺ hematopoietic stem and progenitor cells (HSPCs) were sampled at baseline (day 0) and after 4, 8, and 14 days of differentiation (Fig. 1A,B). In this system, HSPCs reproducibly initiate erythroid, megakaryocytic, basophilic, and myeloid programs with erythroid cells undergoing full terminal maturation and recapitulating all stages of erythropoiesis while other lineages progress through early differentiation stages ^4,45,46^. This system therefore faithfully models early lineage specification and progression, providing a robust framework to investigate mechanisms of cell fate acquisition. Cells from each time point were stained with a panel of 158 barcoded antibodies targeting specific cell surface proteins, along with isotype controls (Table S1). After staining, cells were permeabilized for the ATAC reaction, partitioned using a microfluidic system, and subjected to library preparation and sequencing, following the TEAseq protocol ^29^ (Methods). For integrated data analysis, we used the deep generative model MultiVI (Multi-Variational Inference) ^47^, which employs probabilistic methods to learn a shared latent space that captures cellular states by leveraging all three modalities while accounting for technical effects like batch variation. In this way, all three modalities contribute to defining cell identities. A unique feature of MultiVI is that it explicitly models protein data and corrects for the background noise typical of antibody-based measurements ^47,48^. Notably, MultiVI’s performance has been previously validated on TEA-seq datasets ^47^. Following MultiVI analysis, we obtained a trimodal latent space, which we further projected onto a two-dimensional ForceAtlas (FA) layout ^49^ for visualization. Examination of the FA map reveals that, when the three modalities (RNA, chromatin and cell surface proteins) are integrated, cells are organized as a continuum rather than discrete populations (Fig. 1C).

**Figure 1.**
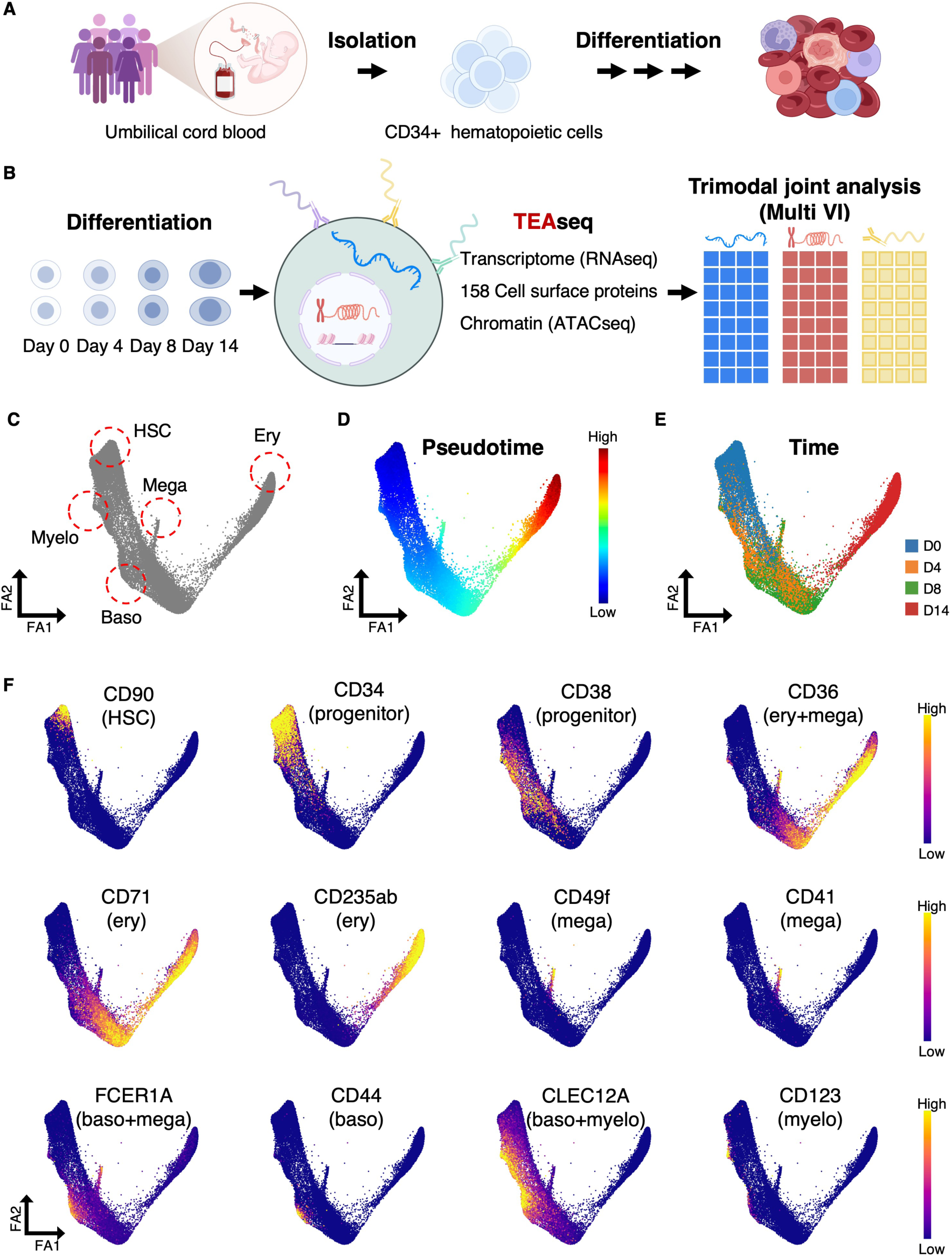
Trimodal Profiling of Human Early Hematopoietic Differentiation. (A) CD34⁺ HSPCs were isolated from umbilical cord blood and differentiated ex vivo. (B) Workflow schematic. Cells collected at four time points were processed with TEA-seq and jointly analyzed with the deep generative model MultiVI. (C) FA map of the MultiVI trimodal latent space (23,887 cells), showing the start cell population (HSC) and the four terminal states identified from a forward Markov chain model of differentiation: myeloid (Myelo), megakaryocyte (Mega), basophil (Baso) and erythroid (Ery). (D) FA map colored by diffusion pseudotime. (E) FA map colored by experimental time points. (F) FA map colored by TotalVI protein expression for 12 marker proteins. See also Figure S1 and Table S1.

To define trajectories, we assigned each cell a pseudotime value and constructed a transport map, modeling forward differentiation as a Markov chain using CD34^+^CD90^+^ hematopoietic stem cells (HSC) as the starting point ^50,51^ (Methods). From the transport map, four terminal states were identified, which we named based on known cell surface proteins ^52–55^: 1) CD123^+^CLEC12A^+^ early myeloid progenitors (**Myelo**), 2) CD49f^+^CD41^+^ megakaryocyte progenitors (**Mega**), 3) CD44^+^FCER1A^+^CLEC12A^+^ basophil progenitors (**Baso**) and 4) CD36^+^CD71^+^CD235ab^+^ differentiated erythroid cells (**Ery**) (Fig. 1C,F). Importantly, the inferred pseudotime ordering (Fig. 1D) aligns well with our experimental time course (Fig. 1E). As expected, both HSPCs and terminally differentiated erythroid cells are predominantly in the G1 phase of the cell cycle, whereas progenitors proliferate extensively (Fig. S1). Thus, this trimodal map recapitulates known features of human hematopoiesis including its gradual nature.

### Establishing a Trimodal-based High-Resolution Map of Human Early Hematopoiesis and Erythropoiesis

To better understand how cells progress along differentiation trajectories, we first estimated the probability of each cell to reach one of the terminal states (Myelo, Baso, Mega, or Ery) using MIRA ^51^, a probabilistic framework that models transcriptional dynamics from multimodal data (Methods) (Fig. 2A, left). Based on these fate probabilities, we reconstructed a lineage tree that arranges cells according to changes in fate potential along pseudotime (Methods) (Fig. 2A, right). This tree-like structure confirms the hierarchical organization of hematopoietic lineages, including separation between the myeloid/basophil and megakaryocyte/erythroid branches, as well as separation between megakaryocyte and erythroid lineages ^8^. Notably, the observed association between basophils and erythroid cells aligns with previous findings ^3,56,57^, including our own single-cell proteomic analyses ^4^. It is important to note that these bifurcations represent biases toward certain lineages rather than definitive fate commitments.

**Figure 2.**
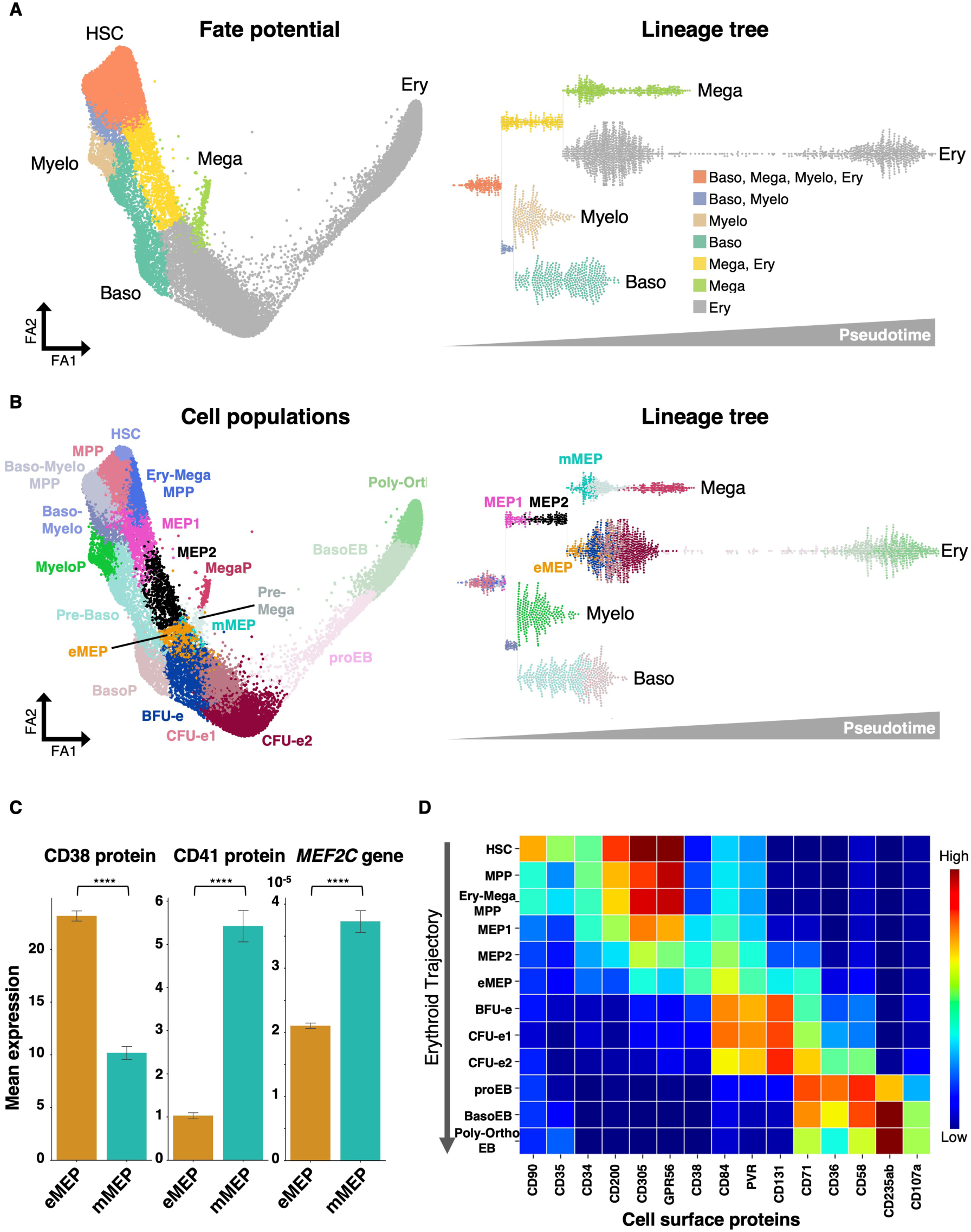
Trimodal-based High-Resolution Map of Human Early Hematopoiesis and Erythropoiesis (A) FA map of the MultiVI trimodal latent space (left) and bifurcating tree arranged as a swarm plot (right), both colored by parsed lineage probabilities. Bifurcations represent lineage biases (Methods) and their ordering reflects pseudotime progression. (B) FA map of the MultiVI latent space (left) and bifurcating tree (right), both colored by cell types defined by Leiden clustering and refined by re-clustering based on known cell type markers. (C) Bar graph showing mean expression of the indicated proteins and genes in defined populations. Data represent mean ± SD. Statistical analysis was performed using an unpaired two-tailed t-test with Welch’s correction. ****p≤0.0001. (D) Heatmap of selected cell surface proteins expression across populations along the erythroid trajectory. X-axis: surface proteins. Y-axis: populations defined in (B). Color scale: TotalVI normalized protein abundance, from low (blue) to high (red). See also Figures S2 and S3 and Tables S2, S3 and S4.

To relate these trajectories to previously defined cell populations, we applied Leiden clustering ^58^, and then refined the clusters by merging or sub-clustering based on both surface proteins (Fig. S2A, Table S2) and transcripts (Fig. S2B, Table S3). See Table S4 for all differentially expressed genes and proteins. This approach yielded a series of distinct populations along hematopoietic trajectories that align well with previously described cell types (Fig. 2B, left). For example, we identified hematopoietic stem cells (**HSCs**; CD34⁺CD38⁻CD90⁺), which expressed MECOM, PRDM16, HLF, and ERG and were non-proliferating. We also found erythroid progenitors (**CFU-es**; CD34⁻CD36⁺CD71⁺CD235ab⁻) characterized by MYC and E2F1 expression and high proliferative activity, as well as terminally differentiated erythroid cells (**Poly-OrthoEBs**; CD36^low^CD71⁺CD235ab⁺) expressing NFE2, TAL1, GATA1, and HBB with low proliferation. In addition, we identified megakaryocyte progenitors (**MegaP**; CD34⁻CD36⁺CD41⁺CD49f⁺) expressing FLI1, MEF2C, and MEIS1, and basophil progenitors (**BasoP**; CD34⁻CLEC12A⁺FCER1A⁺CD44⁺) expressing GATA2, along with many other populations (Fig. 2B; Fig. S2; Fig. S3A; Tables S2–S4).

To refine population identities, we projected them onto the fate probability tree (Fig. 2B, right), which provided a clearer view of lineages and improved separation of certain populations. For example, based on surface marker expression (CD34⁺ CD38⁺ CD36⁻ CD71⁺ CD123⁻) and transcript levels (*GATA1, TAL1, MEIS1, KLF1, FLI1, MYB*), four clusters were identified as megakaryocyte-erythroid progenitors (**MEPs**) ^53,59–61^: pink, black, orange, and turquoise (Fig. 2B). While on the FA map, pink (MEP1) and black (MEP2) can be distinguished based on pseudotime, orange and turquoise clusters overlap (Fig. 2B, left). In contrast, on the lineage tree, they are clearly separated with the turquoise cluster branching toward the megakaryocytic trajectory and the orange cluster branching toward the erythroid trajectory (Fig. 2B, right). We therefore designated these as mMEP (megakaryocyte-biased MEP) and eMEP (erythroid-biased MEP), respectively. Importantly, these assignments are supported by prior literature with the eMEP population expressing higher levels of CD38 (Fig. 2C), consistent with previously validated erythroid-biased MEPs ^59^, whereas the mMEP population expresses higher levels of CD41 and *MEF2C* (Fig. 2C), consistent with previously validated megakaryocyte-biased MEPs ^60,62,63^. These results demonstrate that our integrated approach can distinguish even closely related progenitor states with distinct lineage biases.

Taken together, our analysis resolved multiple cell populations spanning four hematopoietic trajectories. The erythroid branch progresses from HSCs through successive intermediates (including eMEPs, BFU-e, and CFU-e) to terminal Poly-OrthoEBs, while the megakaryocyte branch shares early intermediates but diverges through mMEPs to MegaPs. Parallel basophil and myeloid branches likewise proceed through distinct progenitors, ending at BasoP and MyeloP, respectively (Fig. 2B). Importantly, surface protein profiling along these trajectories not only confirmed gradual phenotypic transitions but also revealed candidate markers, such as GPR56 in HSCs (Fig. 2D; Fig. S3B), which is linked to the maintenance of stemness ^64^.

In summary, by integrating fate probability analysis with multimodal profiling, our study establishes one of the most refined and comprehensive maps of hematopoietic differentiation to date. This framework not only recapitulates well-established lineage relationships but also links the gradual acquisition of hematopoietic fates with specific combinations of known and novel cell-surface markers. All datasets and maps are available for interactive exploration via our companion web application. https://hematopoiesis.systemsbiology.net

### Multimodal Discrepancies Between Protein, RNA, and Chromatin

While protein abundance is often assumed to correlate with transcript levels and chromatin accessibility, whether these relationships consistently hold remains unclear. Our trimodal trajectories provide a unique opportunity to directly contrast these modalities. To begin, we examined representative genes (CD34, CD38, and CD235a) across all three modalities (Fig. 3A). We found that CD235a (glycophorin A) shows strong agreement between protein expression, RNA levels, and promoter accessibility. In contrast, CD38 exhibits promoter accessibility from the earliest stages (HSCs), well before detectable transcript or protein, and CD34 protein persisted far longer than transcript, with promoter accessibility extending even further into erythroid progenitors.

**Figure 3.**
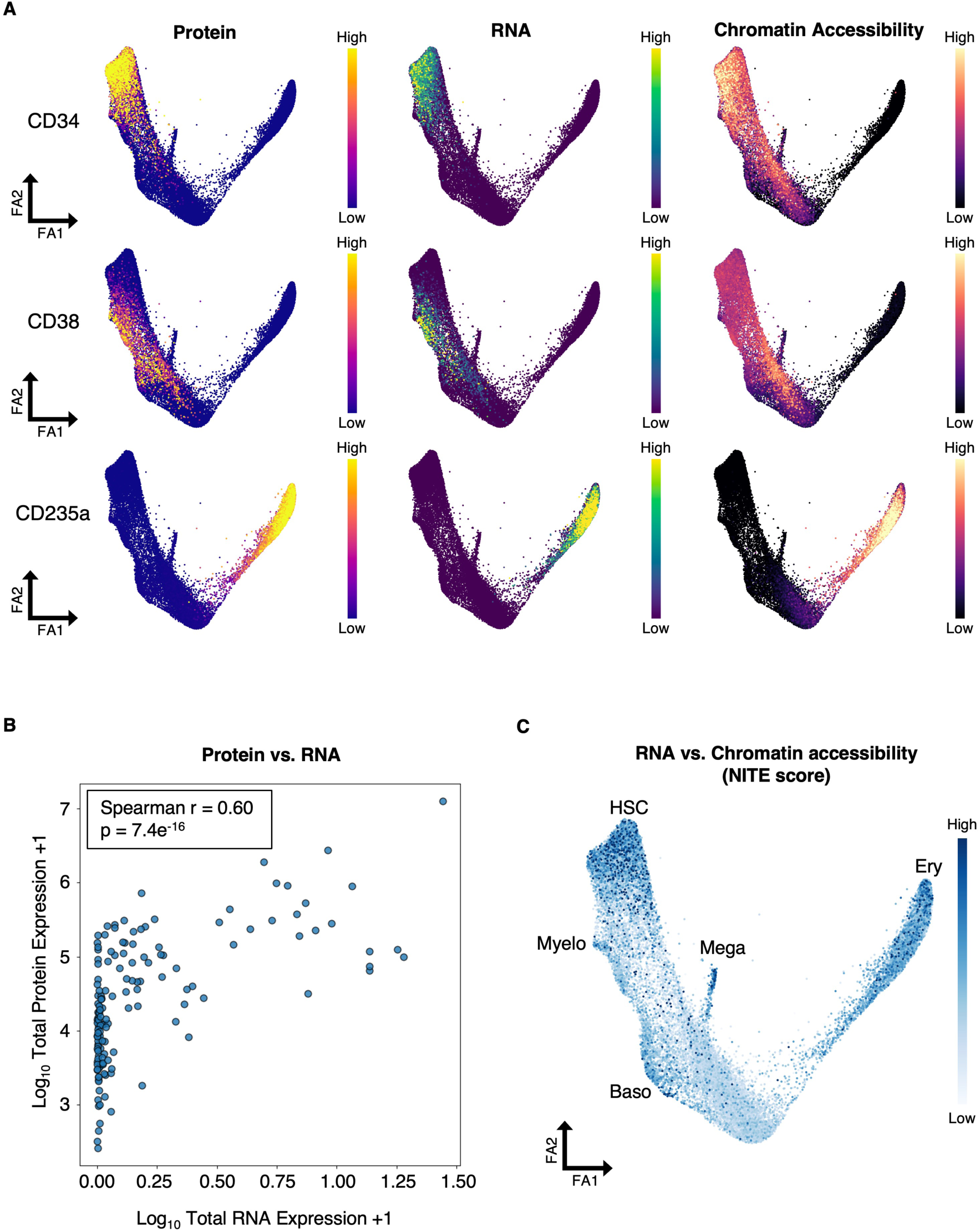
Multimodal Discrepancies Between Protein, RNA, and Chromatin. (A) FA map of the MultiVI latent space colored by TotalVI protein abundance (left), RNA expression (middle), and chromatin accessibility at the promoter (right). (B) Correlation between transcript and surface protein levels across cells. (C) FA map of the MultiVI latent space colored by the NITE score, a per-cell metric that quantifies the divergence between observed gene expression and gene expression predicted from local chromatin accessibility (Methods).

We next quantified these relationships systematically. Correlation analysis between surface protein and transcript levels yielded a moderate correlation (Spearman r = 0.60) (Fig. 3B), consistent with our prior mass spectrometry results^19^. Importantly, many surface proteins remain abundant despite loss of transcripts, underscoring the importance of direct protein measurement rather than inference from mRNA.

To systematically explore the relationship between chromatin accessibility and gene expression, we performed Regulatory Potential (RP) modeling, which predicts gene expression from local chromatin accessibility at promoters and enhancers ^51^ (Methods). We then calculated a <u>n</u>on-locally influenced transcriptional <u>e</u>xpression (NITE) score, a per-cell metric that quantifies how much a gene’s observed expression diverges from the level predicted by local chromatin accessibility ^51^ (Methods). For example, a low NITE score means there is good correlation between chromatin accessibility and gene expression. We found that progenitors in mid-differentiation display the lowest NITE scores, indicating transcription well explained by local chromatin accessibility (Fig. 3C). By contrast, early HSPCs and terminal cells exhibit higher NITE scores, indicating that at these stages, gene expression is less predictable from chromatin accessibility alone and may be more strongly influenced by transcription factor abundance and/or signaling inputs. Together, these results reveal stage-specific discrepancies between modalities.

### Contrasting Chromatin Accessibility and Transcription Reveals Three Phases of Erythropoiesis

To further dissect the interplay between gene expression and chromatin accessibility, we focused on the erythroid trajectory, spanning CD90⁺CD34⁺ HSCs to CD71⁺CD235ab⁺ orthochromatic erythroblasts (orthoEBs) (Fig. 4A, left to right). The strongest divergence between accessibility and transcription (high NITE scores) occurs in multipotent HSCs and MPPs where chromatin is often open before transcription ^20–25^. As cells progress into fate-restricted MEPs and early erythroid progenitors (BFU-e and CFU-e), accessibility and expression become tightly correlated (low NITE scores). Strikingly, the transition from CFU-e to proEBs reverses this trend: accessibility and transcription decouple again, coinciding with a sharp rise in the expression of master regulators such as GATA1, KLF1, and TAL1 (Fig. 4A).

**Figure 4.**
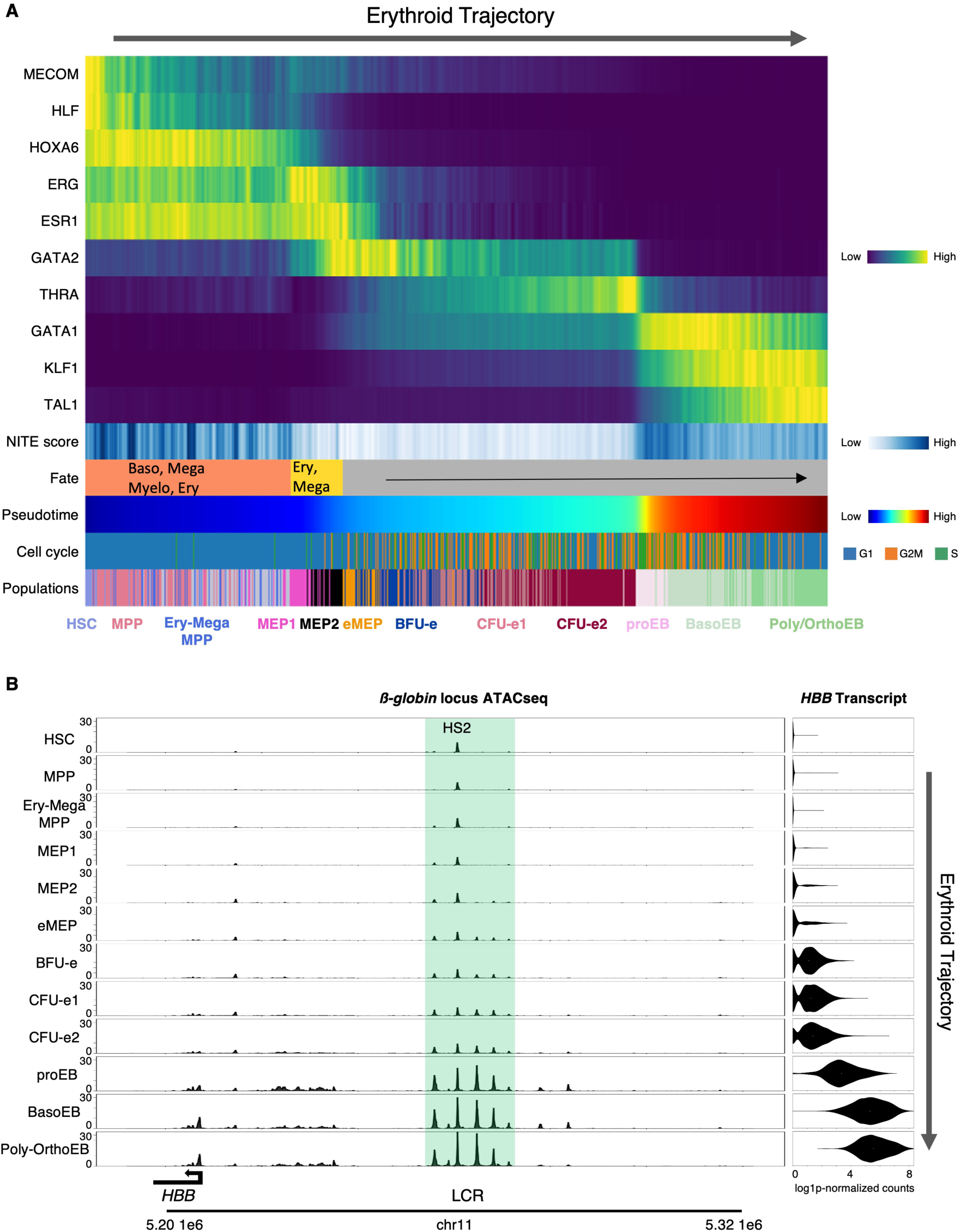
Contrasting Chromatin Accessibility and Transcription Reveals Three Phases of Erythropoiesis (A) Heat map of the erythroid trajectory (left to right) showing: transcript levels for the indicated genes, NITE scores quantifying the divergence between observed gene expression and gene expression predicted from local chromatin accessibility (Methods), lineage biases representing fate potential (as in Fig. 2A), pseudotime (as in Fig. 1D), cell-cycle status (Fig. S1), and populations defined in Fig. 2B. (B) Gradual opening of chromatin at the β-globin locus control region (LCR) coincides with a progressive increase in β-globin transcription along the erythroid trajectory. Chromatin accessibility is shown as ATAC-seq tracks (left), and transcript levels as violin plots (right), in the indicated cell types (defined in Fig. 2B). HS (hypersensitive site). Tracks were downloaded from our web application (https://hematopoiesis.systemsbiology.net).

Collectively, our results reveal three distinct phases of regulatory control along the erythroid trajectory: (1) TF abundance dictates gene expression in early HSPCs; (2) chromatin accessibility governs expression in intermediate progenitors; and (3) during terminal differentiation, TF fluctuations dominate within a chromatin landscape that is already established.

### Complex Changes in Enhancer Activity along Hematopoietic Trajectories

To dissect the relationship between chromatin accessibility and transcription at the gene level, we examined the β-globin locus, a classic model in erythropoiesis. This locus contains five β-like globin genes (e.g., *HBB*) regulated by the locus control region (LCR) where five DNaseI hypersensitive sites (HS1-HS5) mark a super enhancer, which ensures developmental and stage-specific expression ^65,66^. Our TEAseq data show that HS2 is already accessible in HSCs despite absent *β-globin* expression (Fig. 4B). At this stage, however, only HS2 is open. Other LCR enhancers remain closed until late MEPs, when they begin to open alongside the initiation of low-level *β-globin* transcription. Terminal erythropoiesis is then marked by a sharp increase in accessibility across both the LCR and β-globin promoters, coinciding with dramatic upregulation of β-globin expression (Fig. 4B). Thus, enhancer accessibility at this locus progressively increases along the trajectory and correlates closely with transcription.

To test whether such dynamics generalize beyond β-globin, we next examined GATA2, a transcription factor essential in HSCs, MEPs, and basophil progenitors ^67–69^. As expected, its expression fluctuates across trajectories: low in HSCs, transient in MEPs, downregulated in erythropoiesis, and strongly upregulated in basophils (Fig. 5). Enhancer accessibility, however, does not always mirror expression. The intron 5 enhancer (“+9.5 kb” in mouse) is open in HSCs, consistent with its essential role in HSC emergence ^70,71^, but closes along the basophil trajectory despite rising GATA2 expression (Fig. 5B). Conversely, the distal hematopoietic enhancer (G2DHE), ∼116 kb upstream of the TSS (∼77 kb in mouse) ^72–75^, opens gradually in basophil progenitors (Fig. 5B). In MEPs, both enhancers are partially open (Fig. 5A), consistent with genetic evidence that both are required for MEP function ^68^.

**Figure 5.**
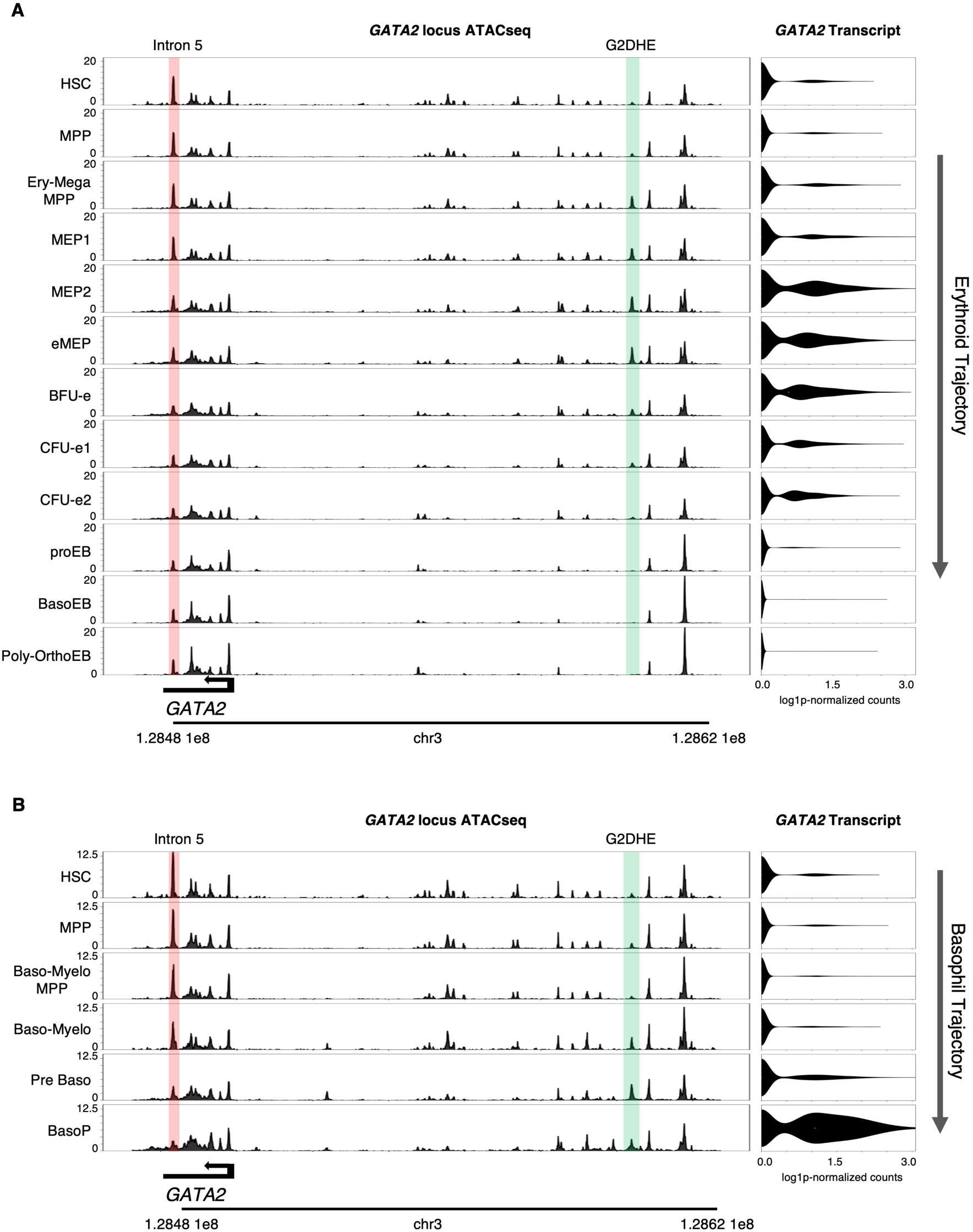
Complex Changes in Enhancer Activity at the GATA2 Locus along Hematopoietic Trajectories (A) Along the erythroid trajectory, the “intron 5” enhancer (highlighted in pink) gradually decreases, while the distal hematopoietic enhancer (G2DHE, highlighted in green) shows transient opening in MEPs. Chromatin accessibility is shown as ATAC-seq tracks (left), and transcript levels as violin plots (right), in the indicated cell types (defined in Fig. 2B). (B) Along the basophil trajectory, the “intron 5” enhancer (highlighted in pink) progressively closes, whereas the G2DHE (highlighted in green) gradually opens. Chromatin accessibility is shown as ATAC-seq tracks (left), and transcript levels as violin plots (right), in the indicated cell types (defined in Fig. 2B). Tracks were downloaded from our web application (https://hematopoiesis.systemsbiology.net). See also Figure S4.

Comparable context-dependent enhancer logic is evident at FLI1, a factor required in HSCs and megakaryocyte progenitors ^4,76–79^, where distinct enhancer elements are differentially accessible across these lineages (Fig. S4).

Taken together, these findings reveal that enhancer regulation along hematopoietic trajectories is highly context- and dose-dependent: some enhancers track transcription closely, others diverge, and their combined activities ultimately determine lineage-specific expression. Thus, enhancer dynamics reveal a regulatory complexity that cannot be captured by looking at chromatin accessibility or transcription alone. They instead call for an integrated approach (enhancer-based gene regulatory networks (eGRNs) ^40,41^) to connect transcription factor activity, enhancer accessibility, and target gene expression into coherent regulatory programs.

### Enhancer-based GRNs Capture Hematopoietic Populations

To better understand how enhancers orchestrate gene programs that control cell fate decisions, we applied SCENIC+, a GRN inference framework that integrates single-cell data on gene expression, chromatin accessibility, motif enrichment, and enhancer-gene associations ^80^. Enhancer-gene links are prioritized using a gradient boosting-based machine learning model, producing enhancer-based regulons (eRegulons) composed of an effector TF, the enhancers and promoters it binds (Regions (R)), and its predicted target genes (G) (Fig. 6A). The activity of each eRegulon in each cell type is defined jointly by TF target genes expression (blue to red color scale) and chromatin accessibility at *cis*-regulatory elements (black dot with varying size) (Fig. 6A). Applying SCENIC+ to our datasets revealed multiple eRegulons with distinct activity patterns across hematopoietic populations, thereby delineating cell-specific eGRNs along lineage trajectories (Fig. 6B).

**Figure 6.**
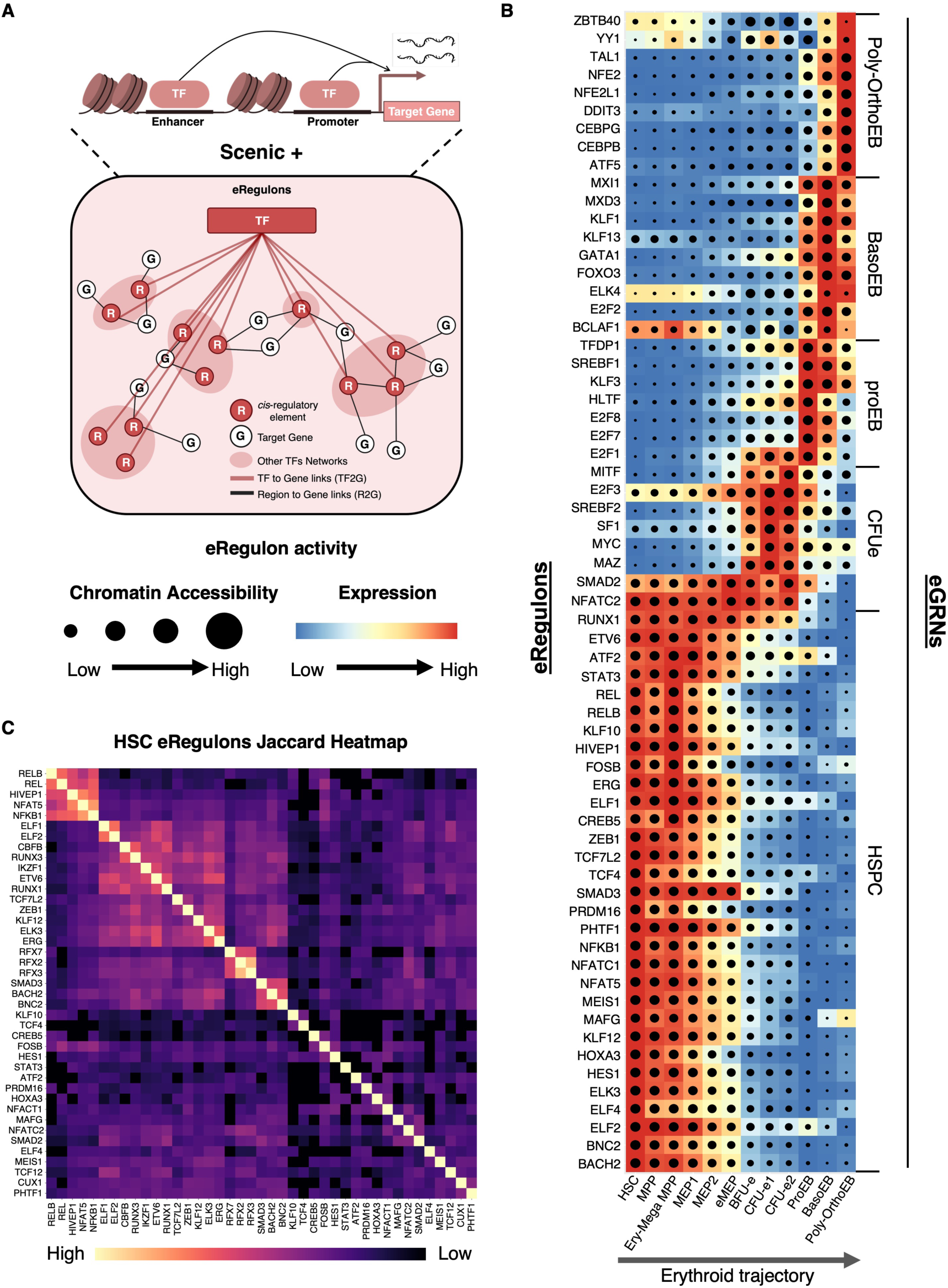
Rewiring of Cell-Specific eGRNs along the Erythroid Trajectory (A) Schematic of eRegulons generated with SCENIC+. (B) Heatmap / dot-plot of eRegulon activity generated with SCENIC+. Each eRegulon consists of an effector TF (listed on the left), the CREs (enhancers and promoters) bound by that TF, and its predicted target genes. Target gene expression is represented by color intensity, and the fraction of CREs with open chromatin is indicated by dot size. Populations (x-axis) are ordered along the erythroid trajectory (as defined in Fig. 2B). Selected eRegulons are shown on the y-axis (left), grouped into eGRNs (right). (B) Heatmap showing target gene overlap among eRegulons. Degree of sharing is represented by the Jaccard similarity coefficient (color scale). Data are available for interactive exploration in our web application (https://hematopoiesis.systemsbiology.net) and in BioTapestry format (Supplemental Data S1). See also Figures S5, S6 and S7 and Table S5.

#### HSPC-specific eGRN

The largest eGRN we identified defines HSPCs and comprises more than 40 eRegulons (29 shown in Fig. 6B) that extensively overlap through shared enhancers and promoters (Fig. 6C). Overlap was quantified using the Jaccard similarity coefficient, which measures the fraction of shared targets between eRegulons (Fig. 6C). Central to this HSPC-specific network are many well-established master regulators of stemness, including ERG ^81–83^, MEIS1 ^84^, ZEB1 ^85^, PRDM16 ^86^, HES1 ^87^, RUNX1 ^88^, ETV6 ^89^, and ELF1 ^90^, among others (Fig. 6, Fig. S5, Fig. S6). To assess the accuracy of this network, we compared predicted TF binding sites with experimentally identified sites in human HSPCs ^91^. This analysis revealed strong overlap for ERG and RUNX1, providing independent validation of our predictions (Fig. S7A,C). Moreover, knockdown of three central regulators (i.e. ERG, RUNX1, and MEIS1) in CD34⁺ HSPCs reduced transcript levels of all tested putative target genes, further validating these as functionally regulated targets (Fig. S7B,D,E). Together, these results provide strong experimental support for the HSPC-specific eGRN identified in our analysis.

#### Erythroid-specific eGRNs

In addition to the HSPC-specific eGRN, we identified cell-specific eGRNs along the erythroid trajectory (Fig. 6B). The highly proliferative colony-forming erythroid (CFU-e) population was characterized by an eGRN centered on pro-proliferative TFs such as MYC, whereas the pro-erythroblast (proEB) population displayed an eGRN dominated by cell cycle regulators including E2F8, E2F7, E2F1 and their obligate partner TFDP1. This is consistent with the critical role of cell cycle control in the CFU-e to proEB transition ^92,93^. At later stages, eGRNs were centered on canonical master regulators of erythropoiesis, including GATA1, KLF1 and FOXO3 (most active in BasoEBs) as well as TAL1 and NFE2 (most active in Poly-OrthoEBs) ^91,94^ (Fig. 6B). To validate the erythroid eGRNs, we leveraged a recent genome-wide CRISPR knockout screen that identified genes essential for terminal erythroid differentiation ^95^. Strikingly, over 80% of these genes were captured within our erythroid eGRNs, confirming that the networks identify functionally relevant regulators (Fig. S7F). Thus, we integrated key regulators of early and late erythropoiesis into dynamic eGRNs that capture known features of erythroid differentiation and connect master TFs, enhancers, and target genes into coherent regulatory programs.

#### Other cell-type eGRNs

In addition to HSPCs and erythroid trajectories, we also identified an eGRN specific to megakaryocyte progenitors, centered on master regulators of megakaryopoiesis, including FLI1, ETS2, and MEF2C ^62,63,78,79,96^ (Fig. S5A). Although the basophil trajectory in our dataset is arrested at an early stage, we were nevertheless able to identify a cell-specific eGRN for early progenitors, which included the known regulator GATA2 ^69^ (Fig. S5B).

Taken together, our findings recapitulate major known regulators as part of complex eGRNs in which genes are linked to their enhancers. Thus, we have established the first dynamic hematopoietic eGRNs that integrate changes in chromatin accessibility at enhancers and promoters with the expression of effector TFs and their target genes as cells progress along hematopoietic trajectories. These eGRNs can be explored as dynamic representations in our interactive web portal (https://hematopoiesis.systemsbiology.net) or using the BioTapestry desktop application ^97^ (Supplemental Data S1). While space limitations preclude listing all target genes in these representations, the full dataset is provided in Table S5. With these principal eGRNs in place, we next leveraged this framework to dissect the mechanisms governing stemness loss and lineage acquisition.

### Gene Regulatory Networks as Functional Units of Cell Identity

In our analysis, we found that most known hematopoietic populations exhibit distinct eGRNs, consistent with their unique regulatory programs (Fig. 6B). However, we were surprised to discover that some immunophenotypically defined populations do not display an eGRN of their own. Instead, they express attenuated versions of eGRNs active in neighboring populations. For example, MPPs and MEPs show weaker versions of the HSC-specific network while BFU-e and CFU-e2 express variants of the dominant CFU-e1 program (Fig. 6B). Taken together, these findings indicate that eGRN-based analysis provides a mechanistically grounded framework for defining cell populations, one that goes beyond surface marker expression or unsupervised clustering of single-cell data. Thus, we propose that eGRNs, because they constitute functional regulatory units that capture cell identity, provide a powerful new framework and practical tool for defining cell types.

### Loss of Stemness and Acquisition of Lineage Identity are Temporally Separated and Occur through Distinct Mechanisms

The eGRNs we established capture the activity of master regulators of stemness and lineage specification, enabling us to dissect the temporal interplay between chromatin accessibility and transcriptional regulation. To this end, we projected representative eRegulons onto the single-cell lineage tree to visualize their activity across cell populations (Fig. 7A). Each eRegulon’s activity is determined by combining two layers of information (Fig. 7B, top left). The first layer captures the overall expression level of the genes regulated by the transcription factor (Color code). The second layer reflects chromatin accessibility at the regulatory regions of those same genes, including both promoters and enhancers (Circle size). Together, these two components provide an integrated view of both the functional output (gene expression) and the regulatory potential (chromatin accessibility) of the eRegulon. The effector transcription factor is shown separately, highlighting its expression as the central regulator along the trajectory (Fig. 7B, top right).

**Figure 7.**
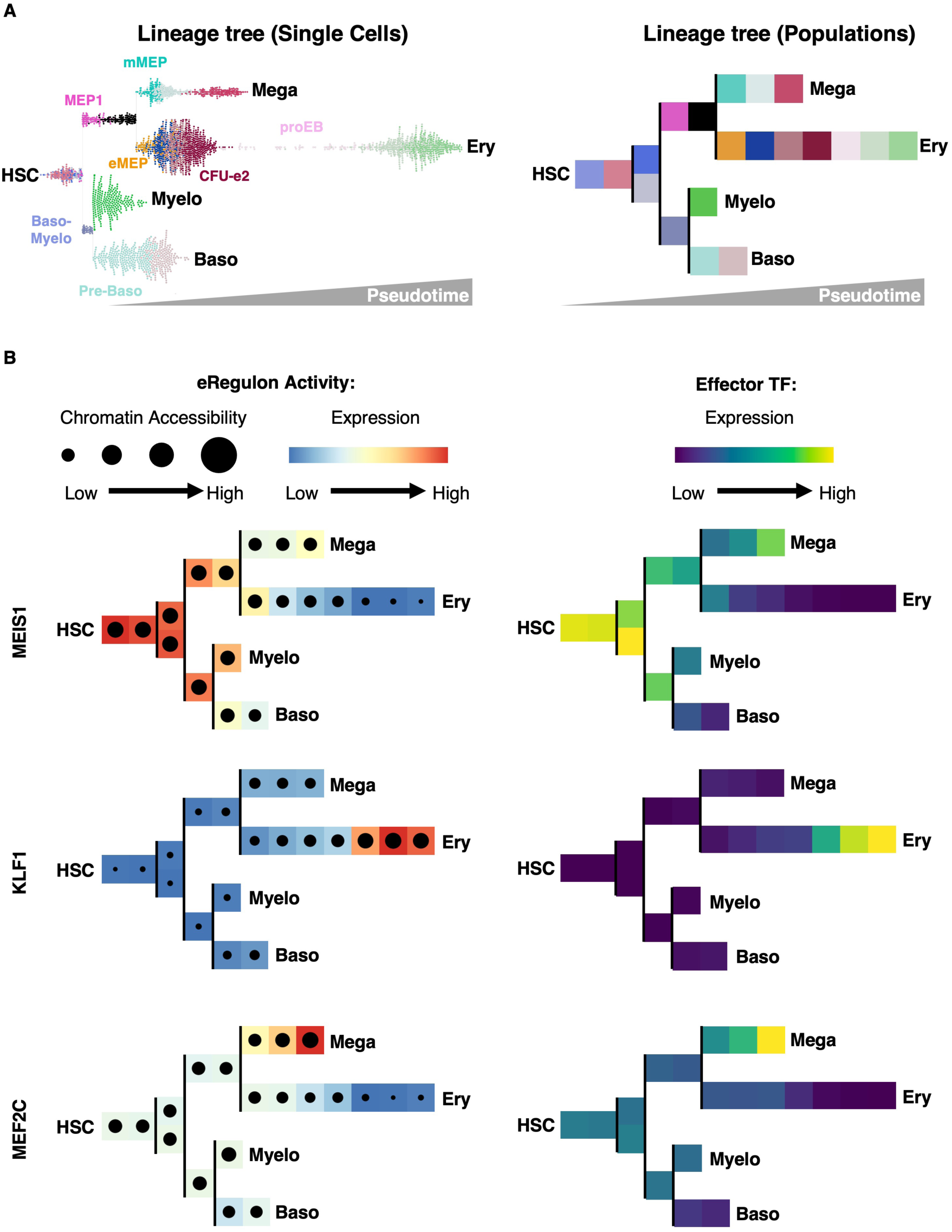
Loss of Stemness and Acquisition of Lineage Identity Are Temporally Separated and Governed by Distinct Mechanisms (A) Left: Bifurcating lineage tree with single cells arranged as a swarm plot colored by cell populations (as defined in Fig. 2B). Right: Bifurcating lineage tree with cell populations (as defined in Fig. 2B). (B) Left: Activity of representative eRegulons for stemness (MEIS1), erythropoiesis (KLF1), and megakaryopoiesis (MEF2C) across hematopoietic lineages shown on the lineage tree. Target gene expression is represented by color intensity, and the fraction of CREs with open chromatin is indicated by dot size. Right: Transcript levels of the corresponding effector TFs across hematopoietic lineages represented on the lineage tree. Expression is represented by color intensity.

To examine stemness loss, we focused on the MEIS1 eRegulon, a central component of the HSC eGRN and an essential regulator of HSC maintenance ^84^. In HSCs, MEIS1 targets are both accessible (large black dot) and highly expressed (red color) (Fig. 7B). Then, their expression progressively decreases across all lineages, yet chromatin at MEIS1 target CREs remains open until late erythropoiesis, closing only in terminal cells (Fig. 7B, left). Thus, in early cells, loss of stemness is driven by reduced MEIS1 abundance (Fig. 7B, right) and diminished transcription of its target genes, rather than chromatin closure that occurs later on the trajectory. This pattern extends to most of the eRegulons in the HSPC eGRN (Fig. 6B). Together, these findings show that stemness loss unfolds in two phases: an initial, reversible transcriptional decline without chromatin closure, followed by late-stage enhancer shutdown by chromatin closing. Notably, this final closure occurs only in advanced progenitors such as CFU-e, underscoring that epigenetic plasticity persists well beyond conventional boundaries of commitment.

We next examined the acquisition of lineage identity, focusing on the KLF1 eRegulon (Fig. 7B, middle left). KLF1 is a key driver of erythropoiesis ^98^ and is central to the BasoEB eGRN (Fig. 6B). We found that in HSCs, the KLF1 regulon is completely inactive, with closed chromatin at its target sites, indicating no priming at this early stage (Fig. 7B, middle left). Instead, priming begins in MEPs, where chromatin at KLF1 targets gradually opens but transcript levels remain low. Several populations later, at the proEB stage, transcription of KLF1 targets sharply increases without additional chromatin remodeling (Fig. 7B). Thus, similar to stemness loss, lineage acquisition proceeds in two phases: a gradual chromatin-opening phase spanning intermediate progenitors, followed by rapid transcriptional activation once effector TF levels rise. This dynamic is also observed for most eRegulons active in terminal erythroid differentiation (proEB, BasoEB, Poly-OrthoEB) (Fig. 6B).

Together, these results reveal that stemness loss and lineage acquisition are governed by temporally and mechanistically distinct processes:

1. Stemness declines first, through reduced TF abundance and transcriptional output in early progenitors.
2. Chromatin remodeling follows, closing stemness enhancers and progressively opening lineage-specific CREs in intermediate progenitors.
3. Lineage identity is then consolidated in terminal progenitors, where high TF abundance drives robust activation of lineage-specific eGRNs within the already remodeled chromatin landscape.

This three-step framework highlights a temporal uncoupling between transcription factor fluctuations and chromatin remodeling, which likely acts as a buffering mechanism to safeguard orderly fate transitions.

### Early Chromatin Priming of Megakaryocyte eGRNs in HSCs

As we found no evidence of erythroid eGRN priming in HSCs, we next asked whether other lineages might behave differently. To test this, we examined eGRN rewiring that underlies the acquisition of megakaryocyte identity, focusing on the MEF2C eRegulon as a representative example. MEF2C plays a critical role in megakaryopoiesis ^62,63^ and is a key component of the eGRN that defines megakaryocyte progenitors (Fig. S5A). In contrast to KLF1, the MEF2C eRegulon is already in an open chromatin state in HSCs, though its target genes show minimal expression, indicating early lineage priming through chromatin accessibility at key enhancers and promoters (Fig. 7, bottom left). This open configuration persists across hematopoietic lineages and closes only during terminal erythroid differentiation. Importantly, MEF2C is not unique: many eRegulons that together form the megakaryocyte eGRN are also chromatin-primed in HSCs (Fig. S5A), in stark contrast to erythroid eRegulons (Fig. 6B). These results reveal an early epigenetic bias of HSCs toward the megakaryocytic lineage but not the erythroid lineage. Together, our findings demonstrate that megakaryocytic eGRNs (unlike erythroid eGRNs) are already chromatin-primed in HSCs, providing a mechanistic basis for the unique ability of HSCs to generate megakaryocytes directly ^99–105^.

## DISCUSSION

In this study, we establish the first dynamic, enhancer-based gene regulatory networks (eGRNs) that resolve the molecular logic of early human hematopoietic fate decisions. Built on a high-resolution, trimodal map of hematopoiesis generated with TEA-seq, these eGRNs integrate transcription factor abundance, enhancer and promoter accessibility, and gene expression within single-cell trajectories anchored to immunophenotypically defined populations. This framework reveals that stemness loss and lineage acquisition are driven by distinct and temporally separated mechanisms: stemness programs decline through reduced TF abundance long before chromatin closure, whereas lineage identity emerges stepwise as enhancers progressively open and lineage-specifying TFs activate their targets. This uncoupling between transcription factor activity and chromatin remodeling provides a buffering mechanism that safeguards orderly and sequential fate transitions. To facilitate the further characterization of genes of interests by members of the community, we deliver a broadly accessible, interactive web portal to explore enhancer-based GRNs and hematopoietic trajectories in unprecedented detail (https://hematopoiesis.systemsbiology.net).

Over the past decade, single-cell assays measuring the transcriptome ^3^, proteome ^4,5^, or chromatin accessibility ^7^ revealed that hematopoietic cells progress along continuous trajectories without sharp boundaries, suggesting fate decisions arise through gradual restriction of potential. Yet, this view conflicts with the well-documented existence of discrete immunophenotypically defined populations that have been isolated and functionally validated ^8–10,53–55^. Even with trimodal profiling of RNA, chromatin, and proteins in the same cells, we observed that each modality still transitioned gradually. Strikingly, however, integrating these modalities into enhancer-based GRNs revealed a very different picture: while stemness is lost gradually through reduced transcription and progressive closure of HSC-specific enhancers, lineage identity emerges stepwise through sequential rewiring of eGRNs by chromatin remodeling and transcriptional activation. This process generates discrete regulatory states that align with immunophenotypically defined populations. Together, these findings reconcile continuous and discrete models of hematopoiesis and establish eGRNs as a powerful framework for defining cell types by their regulatory logic.

Lineage priming refers to the concept that multipotent cells (such as HSCs and MPPs) already possess certain features of terminally differentiated cells, thereby “preparing” them for future activation of lineage-specific programs ^106^. This is important because it suggests that some HSPCs may be biased towards specific lineages if the molecular signatures of one fate are more prevalent ^25^. These ideas originate partly from scRNAseq studies that detected low-level expression of distinct lineage-marker genes (including TFs) in HSCs (transcriptional priming) ^107,108^, and from scATACseq studies that found chromatin opening at CREs of lineage-specific genes prior to their expression (epigenetic priming) ^7,22^. However, the extent to which low-level expression of lineage-specific genes (or the partial opening of their CREs) in HSCs actually reflects functional regulation relevant to lineage bias has remained unresolved. By constructing enhancer-based GRNs that integrate both transcription and chromatin accessibility from single cells, we were able to directly ask whether functional regulatory units that define specific cell types (i.e. eGRNs) are already active at low levels in early multipotential HSCs. Strikingly, when viewed through the lens of eGRNs, we find no evidence of erythroid priming: erythroid-defining eRegulons are completely silent, with closed chromatin and no target gene expression (Fig. 6B). This finding challenges the prevailing assumption that HSCs carry low-level erythroid programs and instead highlights that erythroid identity only emerges later in differentiation (i.e. in fate-restricted MEPs). In sharp contrast, megakaryocyte eGRNs are clearly primed in HSCs (Fig. S5A). Many megakaryocyte eRegulons already display open chromatin and low-level activity, creating a latent regulatory program poised for rapid activation. As cells move along the megakaryocytic branch, these eRegulons become fully activated, whereas along the erythroid trajectory they are progressively extinguished (Fig. 7B). Thus, commitment reflects not only the activation of the chosen program but also the shutdown of competing ones. This early megakaryocytic bias provides a mechanistic explanation for the ability of HSCs to generate megakaryocytes directly, bypassing intermediate progenitors ^99–102^, particularly under stress ^104^ or during aging ^105^. Although not observed in our system, the primed chromatin landscape of megakaryocytic eRegulons makes this potential undeniable (Fig. S5A, Fig. 7B). Together, these findings demonstrate a fundamental asymmetry in lineage priming: while erythroid eGRNs are fully silent in HSCs, megakaryocytic eGRNs are pre-configured and epigenetically poised for activation.

### Limitations of the Study

Our single-cell trimodal study was performed on cord blood–derived HSPCs, which represent early adult hematopoiesis. Prior work has reported functional differences between HSCs from cord blood versus neonatal bone marrow ^109^, underscoring the need to extend these analyses to HSPCs from distinct developmental and anatomical niches. Moreover, our analyses focused on the myeloid, erythroid, and megakaryocytic branches of hematopoiesis; extending this framework to lymphoid differentiation will be an important next step. An additional direction will be to construct enhancer-based GRNs from HSPCs across different ages, particularly in light of recent evidence for an aging-specific direct HSC-to-megakaryocyte route^105^.

## RESOURCE AVAILABILITY

### Lead contact

Request for further information and resources should be directed to and will be fulfilled by the lead contact, Marjorie Brand.

### Material availability

This study did not generate new unique reagents.

### Data and code availability

TEAseq sequencing data have been deposited in the Gene Expression Omnibus (GEO) and are publicly available as of the date of publication.

All trajectories and eGRNs are available for interactive exploration via our companion web application. https://hematopoiesis.systemsbiology.net

This paper does not report original code. Any additional information required to reanalyze the data is available from the lead contact upon request.

## ACKNOWLEDGEMENTS

We thank E. Bresnick (UW-Madison) for critically reading the manuscript. We thank Canadian Blood Services for providing umbilical cord blood samples. We thank S. Boissel and P. Gingras-Gelinas from the IRCM Core Facility (Montreal, QC, Canada) for TEAseq libraries preparation and sequencing. Data analysis was done using the Center For High Throughput Computing (CHTC) at UW-Madison^110^. We thank Brian Bockelman and Christina Koch for their help in navigating the CHTC. This work was supported by NIH (R01-R01DK098449 to M.B and J.A.R), and ASH (Graduate Award to S.T).

## AUTHORS CONTRIBUTIONS

Conceptualization: MB, ST, JAR Methodology: MB, ST, WJRL Formal analysis: ST, SY Investigation: CGP, ST

Funding acquisition: MB, JAR, ST Resources: MB, JAR, RS, FJD Supervision: MB

Visualization: ST, CGP, SY Writing – original draft: MB, FJD

Writing – review & editing: all authors

## SUPPLEMENTAL INFORMATION

**Supplemental Data S1.** Zipped file containing a “README.docx” file with installation instructions for the BioTapestry desktop application, and “AllBTPFiles.zip” file containing cell-specific GRNs in BioTapestry format.

**Table S1**. Antibodies used in TEAseq.

**Table S2**. Differentially expressed surface proteins across populations (lists extracted from MultiVI outputs).

**Table S3**. Differentially expressed genes across populations (lists extracted from MultiVI outputs).

**Table S4**. Full MultiVI differential expression statistics for genes and proteins.

**Table S5**. Enhancer-based gene regulatory networks (constructed with SCENIC+).

**Table S6**. TEAseq quality control metrics.

## SUPPLEMENTARY FIGURES

**Figure S1 (related to Figure 1).**
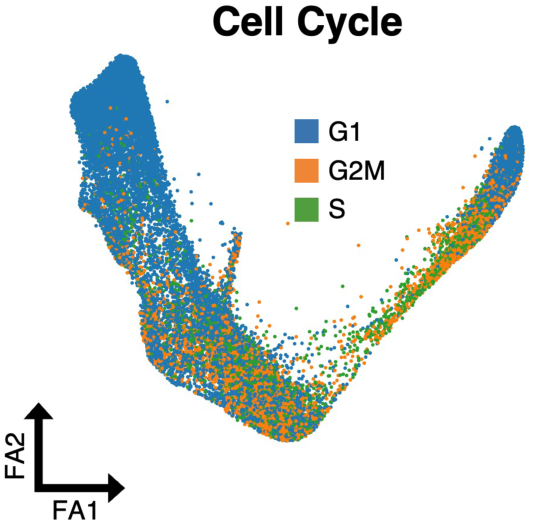
Trimodal Profiling of Human Early Hematopoietic Differentiation. FA map of the MultiVI latent space (23,887 cells) colored by cell cycle phase.

**Figure S2 (related to Figure 2).**
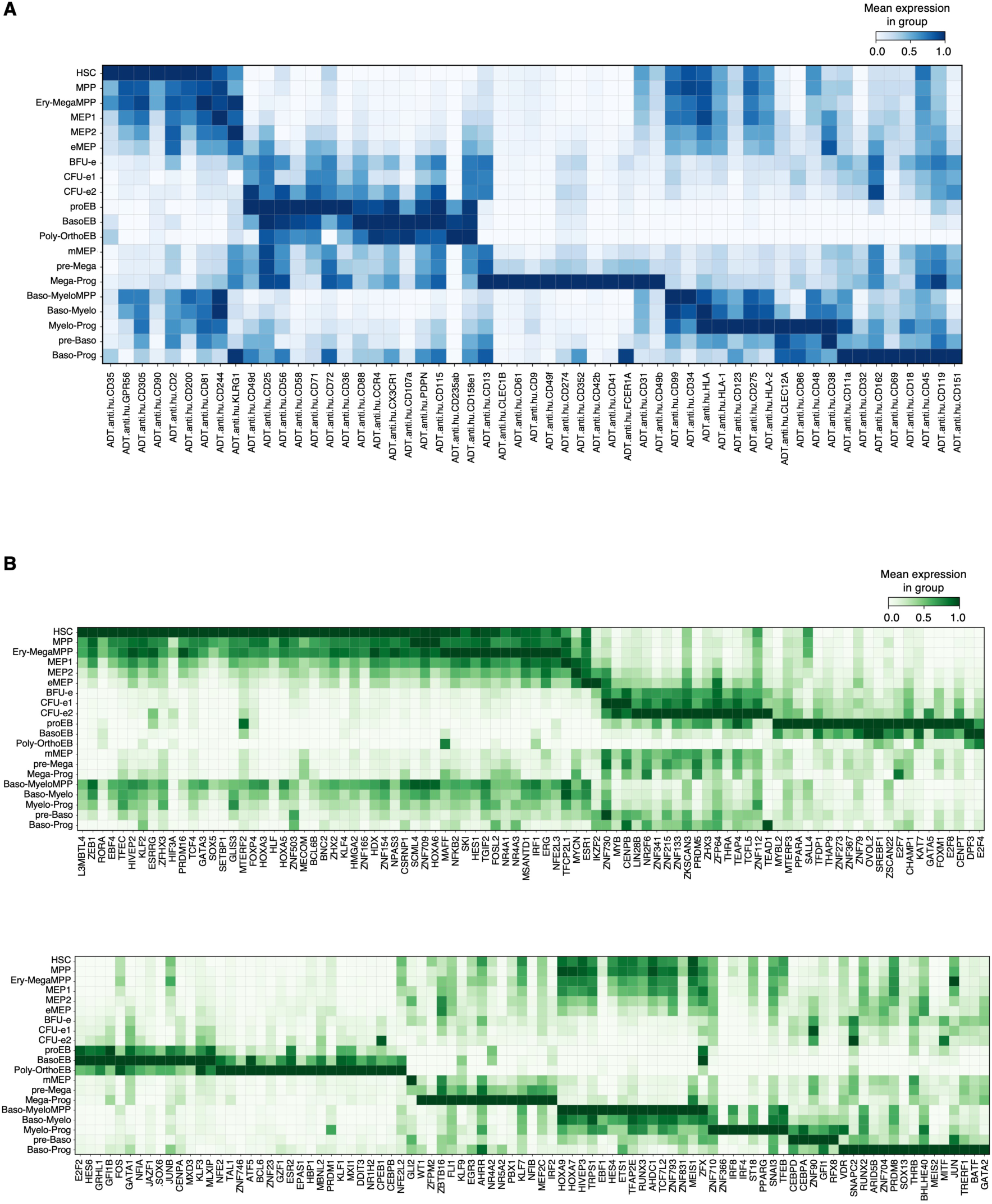
Surface Proteins and RNA Markers of Hematopoietic Populations. (A) Heatmap of selected surface protein markers identified from one-versus-all differential analyses. X-axis: surface proteins; Y-axis: populations defined in Fig. 2B. See Table S2 for the full list of markers. (B) Heatmap of selected RNA markers identified from one-versus-all differential analyses. X-axis: genes; Y-axis: populations defined in Fig. 2B. See Table S3 for the full list of markers. See also Table S4 for the complete results of one-versus-all tests.

**Figure S3 (related to Figure 2).**
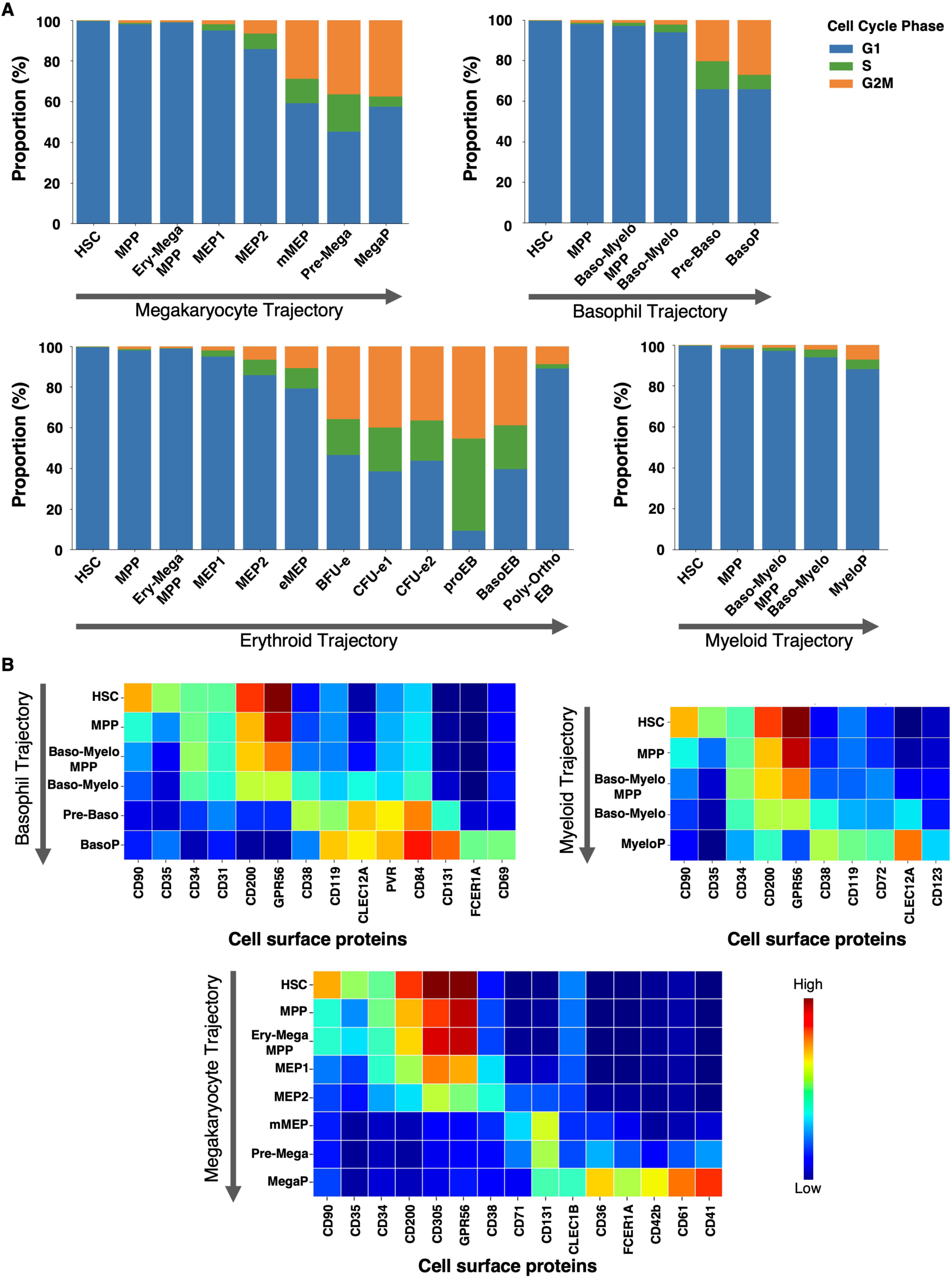
Cell Cycle Distribution and Surface Protein Profiles of Hematopoietic Populations. (A) Bar graphs showing the proportion of cells in each cell-cycle phase across populations along hematopoietic trajectories. (B) Heatmap of selected surface protein expression across populations along the indicated trajectories. X-axis: surface proteins; Y-axis: populations defined in Fig. 2B. Color scale: TotalVI-normalized protein abundance, from low (blue) to high (red).

**Figure S4 (related to Figure 5).**
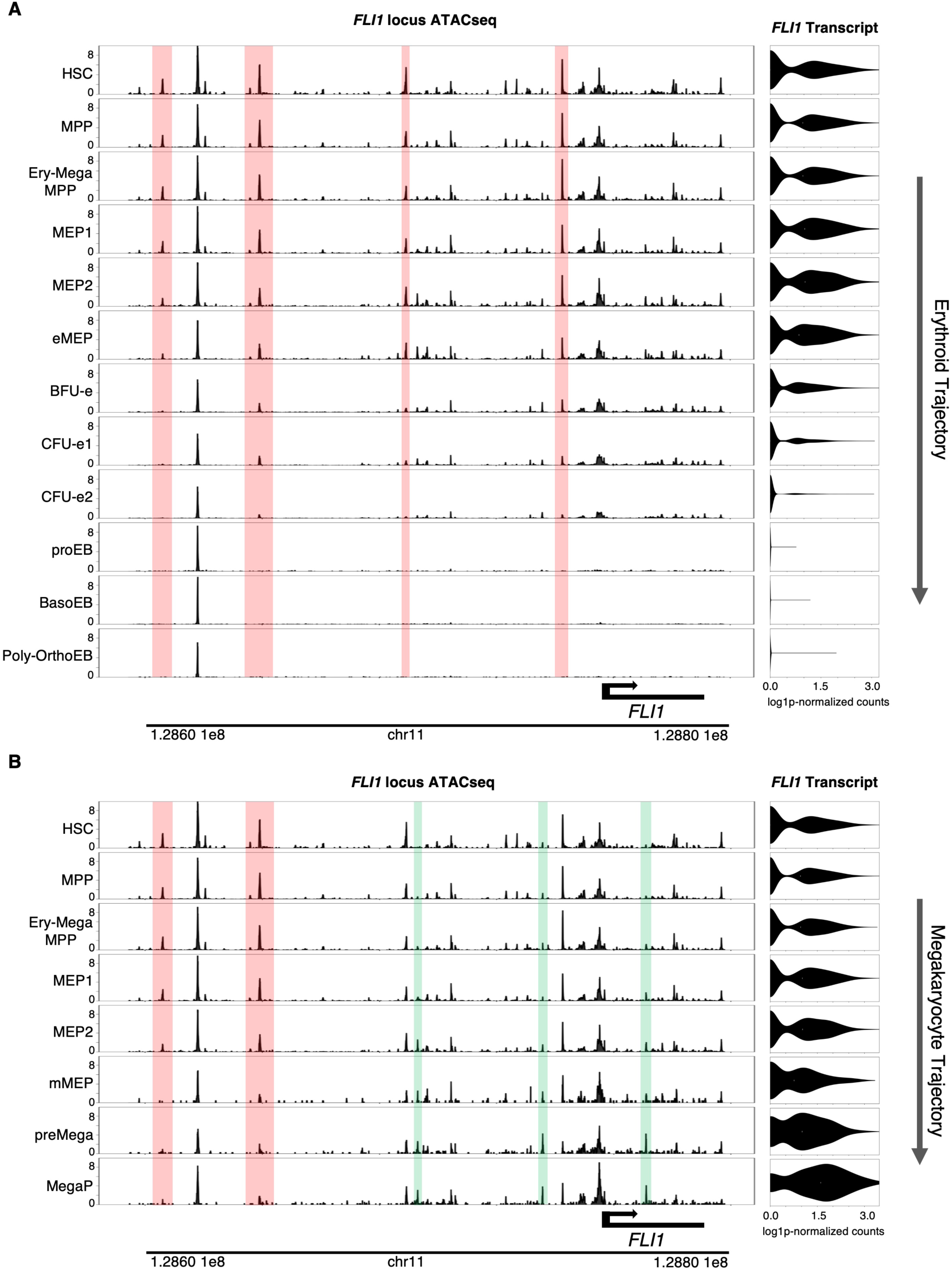
Complex Changes in Enhancer Activity at the FLI1 Locus along Hematopoietic Trajectories (A) Along the erythroid trajectory, progressive decrease in FLI1 expression is accompanied by gradual closure of several enhancers (highlighted in pink). Chromatin accessibility is shown as ATAC-seq tracks (left), and transcript levels as violin plots (right), in the indicated cell types (as defined in Fig. 2B). (B) Along the megakaryocyte trajectory, progressive increase in FLI1 expression is accompanied by gradual opening of some enhancers (highlighted in green) and closure of others (highlighted in pink). Chromatin accessibility is shown as ATAC-seq tracks (left), and transcript levels as violin plots (right), in the indicated cell types (as defined in Fig. 2B). Tracks were downloaded from our web application (https://hematopoiesis.systemsbiology.net).

**Figure S5 (related to Figure 6).**
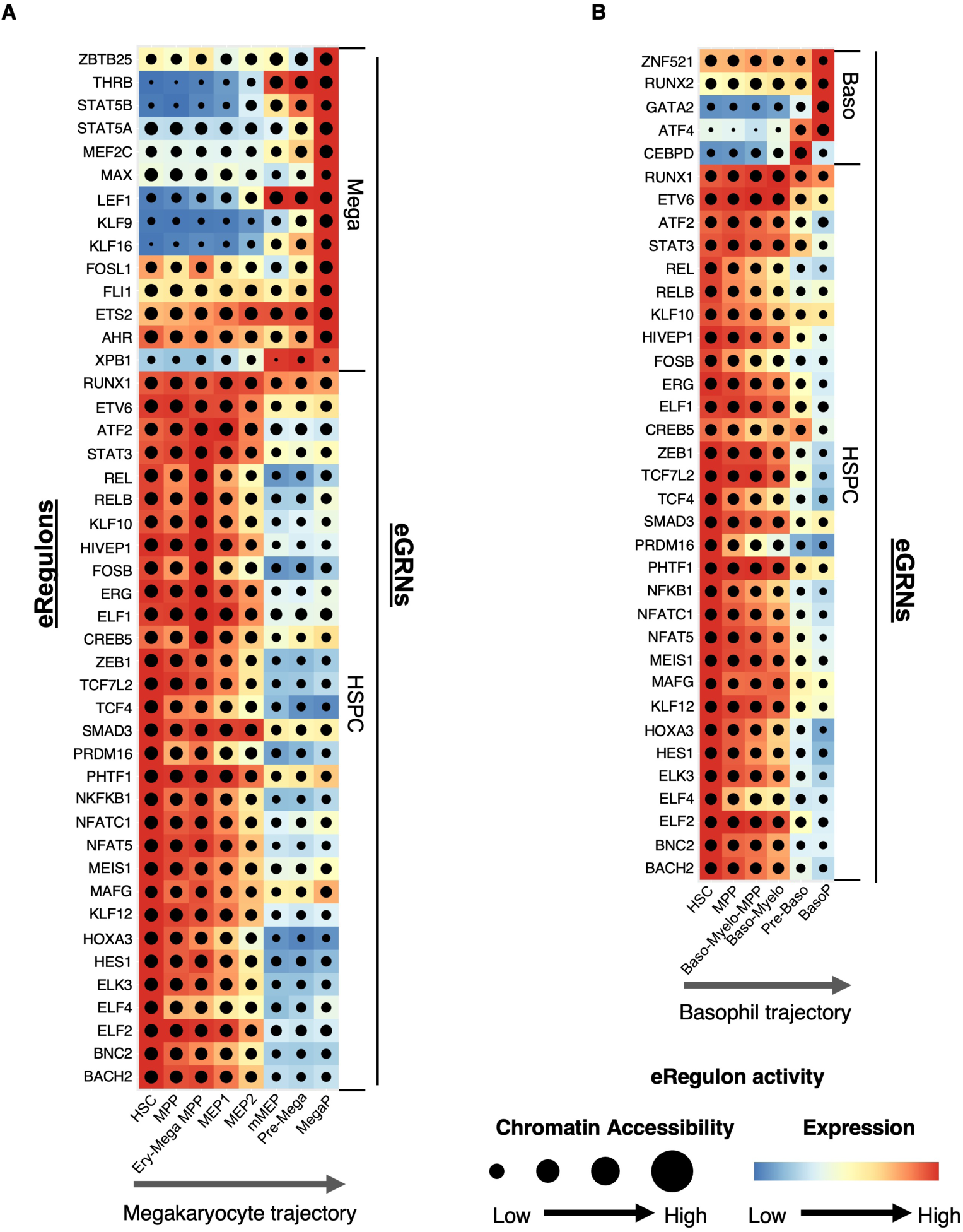
Rewiring of Cell-Specific eGRNs along Hematopoietic Trajectories Heatmap / dot-plot of eRegulon activity generated with SCENIC+. Each eRegulon consists of an effector TF (listed on the left), the CREs (enhancers and promoters) bound by that TF, and its predicted target genes. Target gene expression is indicated by color intensity, and the fraction of CREs with open chromatin is represented by dot size. Populations (x-axis) are ordered along the megakaryocyte (A) and basophil (B) trajectories (as defined in Fig. 2B). Selected eRegulons are shown on the y-axis (left), grouped into eGRNs (right). See Table S5 for the full list of TFs and target genes. Data are available for interactive exploration in our web application (https://hematopoiesis.systemsbiology.net) and in BioTapestry format (Supplemental Data S1).

**Figure S6 (related to Figure 6).**
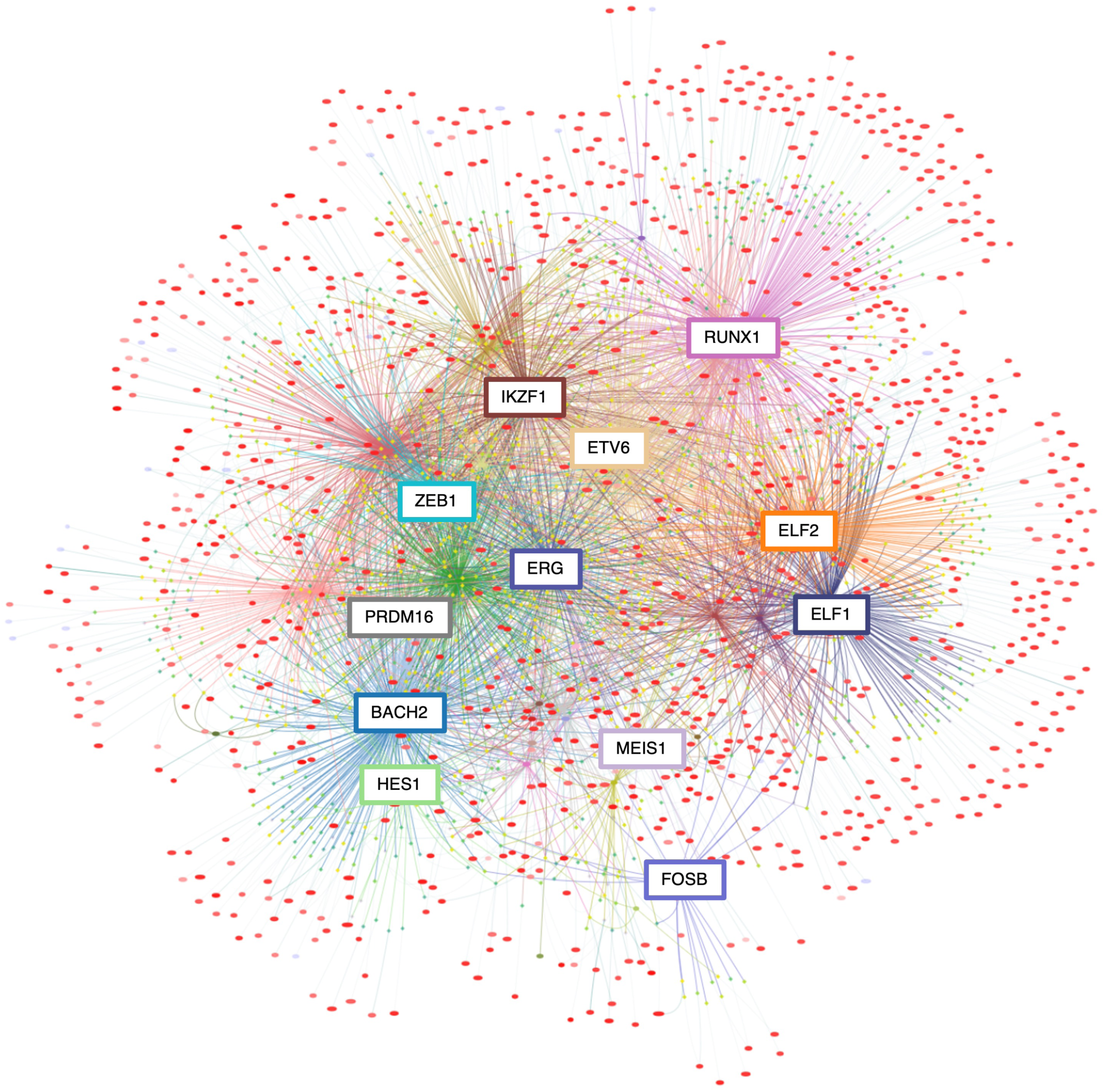
HSPC Enhancer-Based Gene Regulatory Network. Selected eRegulons from the HSPC eGRN are shown. Effector TFs are highlighted in colored boxes. Circles represent target genes (expression shown on a red [high] to blue [low] scale), and diamonds represent cis-regulatory elements (promoters and enhancers; chromatin accessibility shown on a yellow [high] to dark blue [low] scale). For clarity, only the top 4,000 most variable target genes are displayed. See Table S5 for the full list TFs and target genes. The eGRN shown here was retrieved from our interactive web application (https://hematopoiesis.systemsbiology.net). All networks, including this one, are available both online for interactive exploration and as BioTapestry files in Supplemental Data S1.

**Figure S7 (related to Figure 6).**
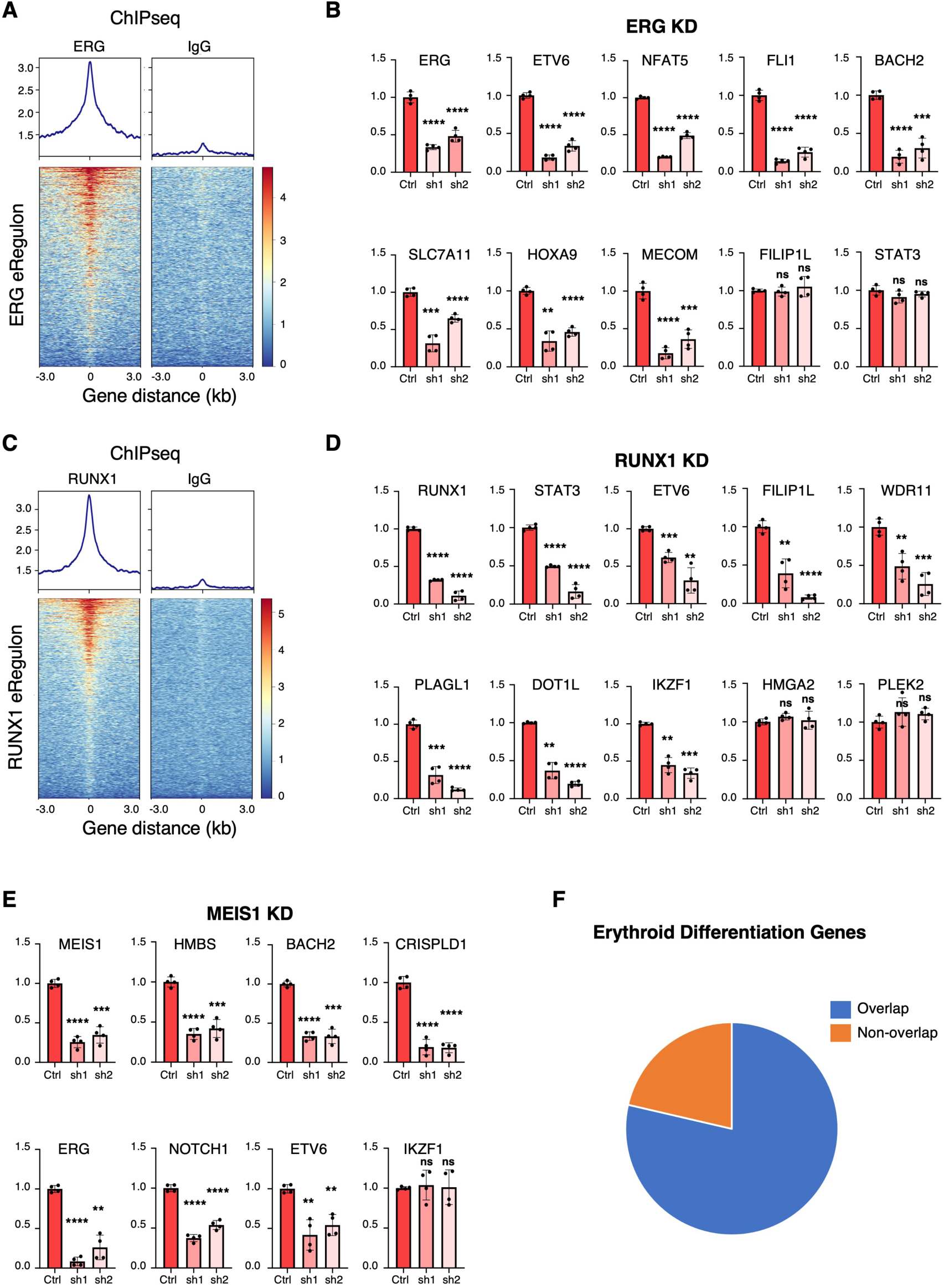
Experimental Validation of Enhancer-Based Gene Regulatory Networks. (A) ERG binds to predicted target CREs within the ERG eRegulon. Published ChIP-seq data ^91^ showing ERG occupancy at predicted sites are displayed as a heatmap; IgG ChIP-seq served as a control. (B) ERG activates its predicted target genes as part of the ERG eRegulon. ERG knockdown (KD) was induced by lentiviral delivery of two shRNAs (sh1, sh2) in CD34⁺ HSPCs. FILIP1L and STAT3 were used as negative controls. (C) RUNX1 binds to predicted target CREs within the RUNX1 eRegulon. Published ChIP-seq data ^91^ showing RUNX1 occupancy at predicted sites are displayed as a heatmap; IgG ChIP-seq served as a control. (D) RUNX1 activates its predicted target genes as part of the RUNX1 eRegulon. KD was induced as in (B). HMGA2 and PLEK2 were used as negative controls. (E) MEIS1 activates its predicted target genes as part of the MEIS1 eRegulon. KD was induced as in (B). IKZF1 was used as a negative control. (F) Overlap between genes essential for erythroid differentiation (identified in a published genome-wide CRISPR screen ^95^) and erythroid-specific eGRNs. (A, C) Signal intensity represents RPKM-normalized ChIP-seq coverage. (B, D, E) Expression of predicted target genes shown by qRT-PCR relative to GAPDH (mean ± SD, n=4). Two-tailed t test: ns, not significant (p > 0.05); *p ≤ 0.05; **p ≤ 0.01; ***p ≤ 0.001; ****p ≤ 0.0001.

## STAR METHODS

### EXPERIMENTAL MODEL

#### Cell acquisition and Ethics

Umbilical cord blood (CB) after healthy, full-term pregnancies was obtained through Canadian Blood Services “Cord Blood for Research program” (CBR-2019-003). All procedures were approved by the Canadian Blood Services Research Ethics Board (REB 2019.027) and the Ottawa Health Science Network Research Ethics Board (REB 2007804-01H).

Murine bone marrow stromal MS-5 cells were obtained from DSMZ (cat#ACC 441; RRID: CVCL_2128) and cultured as follows. Cells are seeded at 2 million cells/80 cm2 (for a 10 cm dish) in alpha-MEM medium containing ribo- and deoxyribonucleosides (Gibco, cat# 12571-048) supplemented with 10% FBS, 2 mM L-glutamine, 2 mM sodium pyruvate, and 1% penicillin/streptomycin. Confluent cultures are split 1:3 every third day using 0.05% Trypsin/EDTA.

Human embryonic kidney 293T cells were obtained from ATCC (cat#CRL-3216; RRID: CVCL_0063) and cultured as follow. Cells are seeded at 1 million cells/80 cm2 (for a 15 cm dish) in Dulbecco’s Modified Eagle’s Medium (DMEM)/High Glucose (HyClone, cat# SH30243.01) supplemented with 10% FBS and 1% penicillin/streptomycin. Confluent cultures are split 1:5 every two to three days using 0.05% Trypsin/EDTA.

## METHOD DETAILS

### Hematopoietic stem and progenitor cells isolation

CD34-positive hematopoietic stem/progenitor cells were isolated as previously described ^4,46^ with some modifications. First, CD34+ cells were enriched from fresh cord blood by negative selection using the RosetteSep Human Cord Blood CD34 Pre-Enrichment Cocktail (STEMCELL Technologies cat#15631), followed by Ficoll density gradient and CD34 positive selection using the Human CD34 MicroBead Kit (Miltenyi, cat#130-046-703) according to the manufacturer’s instructions. Purified cells were analyzed by FACS for CD34 expression using the APC Mouse Anti-Human CD34 antibody (BD Pharmingen, cat# 555824) and either cryopreserved in 10% DMSO or cultured directly as described below.

### Ex vivo hematopoietic culture

CD34-positive hematopoietic stem/progenitor cells were differentiated using a previously described protocol ^4,45,46^. In a first step (day 0 to day 11) CD34+ cells were cultured in serum-free IMDM medium supplemented with 1% penicillin/ streptomycin, 4×10^−3^ M L-glutamine, 40 ug/ml inositol, 10 ug/ml folic acid, 1.6×10^−4^ M monothioglycerol, 90 ng/ml ferrous nitrate, 900 ng/ml ferrous sulfate, 20% albumin-insulin-transferrin (BIT) or differentiation medium (DM), also containing the following cytokines: 10^−6^ M hydrocortisone (HC), 100 ng/ml stem cell factor (SCF), 5 ng/ml interleukin 3 (IL-3) and 3 IU/ml erythropoietin (EPO) for 8 days followed by 3 days in supplemented IMDM medium containing only SCF and EPO. In a second step (day 12 to day 14), cells were co-cultured on a layer of stromal MS-5 cells in the supplemented IMDM medium containing only EPO. Cells were harvested at the indicated Day 0, Day 4, Day 8, and Day 14 time points and cryopreserved in their respective culture media supplemented with 10% DMSO.

### TEAseq procedure

Cells at Day 0, Day 4, Day 8 and Day 14 cells were thawed in complete IMDM for 1 minute at 37°C and centrifuged at 300g for 5 minutes. Supernatant was discarded and cells were resuspended in complete differentiation medium (DM) supplemented with cytokines corresponding to their stage of differentiation, as follow: Day 0 and Day4 in complete IMDM media supplemented with 10^-6^ M Hydrocortisone, SCF (100ng/ml), IL-3 (5ng/ml), and EPO (3UI/ml); Day 8 in complete IMDM media supplemented with SCF (100ng/ml) and EPO (3UI/ml); Day 14 in complete IMDM media with EPO (3UI/ml) only, and incubated for one hour at 37°C. Cells were then centrifuged at 300g for 5 minutes at 4°C, the supernatant discarded, and cells resuspended in 10 ml of ice-cold Cell Staining Buffer (CSB) (BioLegend, cat# 420201), filtered through a 40 μm strainer, and centrifuged at 300g for 5 minutes at 4°C. The supernatant was discarded, and cells were processed for TEAseq as described ^29^ with some modifications. Three million cells for each condition were transferred into 5 mL polystyrene round-bottom tubes and centrifuged at 300g at 4°C for 5 minutes. The supernatant was removed, and cells were resuspended in 22.5 μl of ice-cold CSB with 2.5 μl of TruStain FcX™ PLUS blocking reagent (BioLegend, cat# 422301) and incubated for 5 minutes on ice. Total Seq A antibodies (TotalSeq-A Human Universal Cocktail, V1.0, BioLegend, cat# 399907) supplemented with CD34 (0.125 μg/test for up to 3 million cells), CD90 (1 μg/test for up to 3 million cells), CXCR4 (2 μg/test for up to 3 million cells), and GPA (0.5 μg/test for up to 3 million cells) spiked-in antibodies (Table S1) were prepared following BioLegend guidelines. Briefly, 25 μl of TotalSeqA antibody cocktail was added to the 25 μl of blocked cells, followed by a 30-minute incubation on ice with mixing every 5 minutes. Cells were then diluted with 2 mL of ice-cold CSB, centrifuged, and the supernatant removed. Cells were resuspended in 1 mL of ice-cold CSB, transferred to 1.5 mL low-bind Eppendorf tubes, and centrifuged at 300g for 5 minutes at 4 °C. Wash Buffer was prepared by adding 20 mM Tris-HCl pH 7.4, 150 mM NaCl, and 3 mM MgCl to HyPure Molecular Biology Grade Water (Cytiva, SH30538.03). Cell pellets were resuspended in 100 μl ice-cold Wash Buffer containing 0.01% digitonin (Millipore Sigma, cat# 300410) (Perm Buffer) and incubated for 5 minutes on ice. Cells were then diluted by adding 1 ml of ice-cold Wash Buffer and centrifuged at 300g at 4°C for 5 minutes. Cell pellets were then resuspended in 50 μL of ice-cold Wash Buffer containing 1U/µL Roche’s Protector RNase Inhibitor (Millipore Sigma, cat# 3335399001) (Tagmentation Buffer). Cell suspensions were filtered through a 40 μm strainer, diluted to 3,000 cells per μL, and approximately 6,000 cells were submitted to the next steps for each sample time point. Tagmentation and separation of cells on the Chromium microfluidic system were performed exactly as described ^29^. Libraries were prepared by the IRCM Core Facility (Montreal, Canada) according to the Chromium Next GEM Single Cell Multiome ATAC + Gene Expression User Guide (CG000338 Rev A). Libraries were sequenced at Genome Quebec on an Illumina NovaSeq 6000.

### TEAseq data preprocessing

Data analyses were conducted using the Center for High Throughput Computing (CHTC) platform at the University of Wisconsin-Madison ^110^. For each sample, the RNA and ATAC libraries were aligned to the human GRCh38 reference genome (refdata-cellranger-arc-GRCh38-2020-A-2.0.0) using Cell Ranger ARC v2.0.2. The processing outputs from all four sample time points were aggregated to generate a single feature-barcode matrix using the command line cellranger-arc aggr without sequencing depth downsampled. For Antibody-Derived Tag (ADT) libraries, raw base call files were demultiplexed into FASTQ files using Cell Ranger v7.1.0 and bcl2fastq2 v2.20. For each sample, ADT count matrices were generated using the tag quantification pipeline CITE-seq-Count v1.4.5 available at https://github.com/Hoohm/CITE-seq-Count run with default parameters and the expected number of cells set to 12,000 per sample.

The Cell Ranger ARC filtered RNA and ATAC feature-barcode matrix, and the ADT count matrices were combined into a single MuData object for downstream analyses. Quality control (QC) was performed using Scanpy v1.9.8. QC metrics were calculated without log transformation (log1p=False) for RNA, ATAC, and protein data respectively using Scanpy’s ‘pp.calculate_qc_metrics’ function. Mitochondrial genes were identified as those with gene symbols prefixed by “MT-”. Genes detected in fewer than three cells (RNA), surface proteins detected in fewer than three cells (ADT), and chromatin accessibility peaks present in fewer than 0.5% of cells (ATAC) were excluded. Cells were retained if they expressed more than 350 genes, contained more than 300 peaks, fewer than 5,000 genes, less than 40% mitochondrial reads, and fewer than 5,000 ADT UMIs.

To examine antibody specificity, we applied the totalVI model from scvi-tools (v0.20.3) to jointly model ADT with RNA ^48^. Top 4000 highly variable genes were selected using the Seurat v3 method ^111^ implemented by Scanpy’s ‘pp.highly_variable_genes’ function. The total VI model was trained with a maximum number of epochs set to 400 and default values for all other parameters. Denoised expressions were obtained using the ‘get_normalized_expression’ function from scvi-tools with parameters n_sample=25 and return_mean=True. We assessed totalVI modeled protein foreground probability and denoised protein expression to remove cells exhibiting nonspecific antibody binding or high technical noise. Clusters with elevated expressions for a broad range of proteins in the panel and cells with high foreground probabilities for isotype controls were discarded from the analysis. At the end of the preprocessing, the dataset contained 23,887 cells, 28,431 genes, 165,589 peaks, and 158 proteins (Table S6).

### Trimodal data analyses with MultiVI

Integrated paired analysis of the 3 modalities (single-cell RNA sequencing (scRNA), surface proteins, and single-cell ATAC sequencing (scATAC-seq)) of TEAseq was performed using a pipeline that combines several functions of scvi tools ^112^, multiVI ^47^, totalVI ^48^ and scanpy ^113^. Top 4000 highly variable genes were selected using the Seurat v3 method ^111^ implemented by Scanpy’s ‘pp.highly_variable_genes’ function. The batchkey parameter was set for combined data from all cells, as this data originated from the same TEAseq experiment. We then created the multiVI model using the scvi.model.MULTIVI.setup_anndata function, specifying the name of each layer to be considered, including protein expression, gene expression, and peaks. Protein expression was stored as an obsm attribute to be integrated as a third modality in the model training setup. We left other parameters at default values, including the model training, which was set to the default of 500 epochs. After training, we applied the neighborhood function from Scanpy and used Force Atlas 2 (FA) ^49^ to project the trimodal latent space onto a 2D graphical plot. The Leiden algorithm ^58^ with default resolution was used to generate clusters. We adjusted the resolution and cluster annotations based on the expression of key proteins or genes. The data was stored as an AnnData object for downstream analyses. The AnnData object contained the 23,887 cells, 28,431 genes, 165,589 peaks, and 158 proteins (Table S6).

### Trajectory Analysis with MIRA

We used the preprocessed AnnData object from MultiVI and loaded it for trajectory analysis. Trajectory reconstruction was carried out using MIRA (v2.1.1) ^51^. We used the force-directed FA embedding (X_draw_graph_fa) as the reference layout for visualizations. For pseudotime inference, the starting cell was defined automatically as the cell with the minimum value in the first diffusion map component. Pseudotime ordering was obtained using the get_transport_map function, and results were visualized on FA embeddings. Terminal cell populations were identified with the find_terminal_cells function, run with 3 seeds and 20 iterations to ensure robustness. Both the initial terminal cells and the expanded set (referred to as MORE terminal cells) were projected on the FA graph to confirm biological plausibility. We then computed lineage branch probabilities toward four specified fates: myeloid, megakaryocyte, basophil, and orthoerythroblast. The global tree structure of trajectories was inferred using a threshold of 0.6 and visualized with Scanpy’s draw_graph. To further characterize dynamics, categorical metadata were converted to string types and displayed using MIRA’s plot_stream function. This included lineage progression along pseudotime for merged cell clusters, sample of origin, and MultiVI-derived latent factors. Both swarm-style and probability stream plots were generated to capture changes in cell-type abundance and lineage assignment along pseudotime.

### Differential Expression

Differential expression (DE) analysis was performed using the model.differential_expression function from scvi-tools ^112^. The DE dataframe, de_df, was sorted by the log-fold change median (lfc_median), in descending order, and further filtered to include only genes with a log-fold change median greater than 0.2. Two subsets were created: one for protein (pro) and another for RNA (rna). For protein data, ADT were further filtered to include those with a Bayes factor greater than 0.7. For RNA data, the Bayes factor threshold was set at 1.5, with an additional filter applied to include genes with a non-zero proportion set at 4%. A dendrogram was constructed using Scanpy’s “tl.dendrogram” function, which considers the representation learned by the totalVI (X_totalVI). The results of the differential expression analysis were visualized using a dot plot generated by Scanpy’s “.dotplot” function.

### RNA and ATAC topic modelling with MIRA

We used MIRA v2.1.1 to systematically compare transcription and chromatin accessibility in the same single cells ^51^. All MIRA analyses closely followed the online tutorials.

Topic modeling was first performed on RNA and ATAC separately to generate interpretable latent variables using a variational autoencoder (VAE) neural network. RNA topics represent co-regulated genes, and ATAC topics represent co-accessible genomic loci.

To model RNA topics, raw expression counts were used as input. Top 4000 highly variable genes were selected using the Seurat v3 method ^111^ implemented by Scanpy’s ‘pp.highly_variable_genes’ function. These highly variable genes were included as VAE features and captured in the topics. The expression topic model was instantiated using the ‘make_model’ function. Sample time points (Day 0, Day 4, Day 8, Day 14) were encoded as categorical covariates. The learning rate bounds were set to 1 × 10^-3 and 0.1 to cover the portion of the learning rate versus loss curve with the steepest slope. To estimate a reasonable range for the number of topics, a coarse sweep was first performed with a gradient-based tuner, which suggested an optimal topic number of 20. The model was then fine-tuned using the Bayesian optimization method with a minimum number of topics set to 10, a maximum number of topics set to 35, and default values for other parameters. To model ATAC topics, binarized peak counts were used. All peaks were included as VAE features and captured in the topics. The accessibility topic model was instantiated using the ‘make_model’ function. Sample time points (Day 0, Day 4, Day 8, Day 14) were encoded as categorical covariates. The learning rate bounds were set to 1 × 10^-3 and 0.1 to cover the portion of the learning rate versus loss curve with the steepest slope. The model was tuned using the Bayesian optimization method with a minimum number of topics set to 11, a maximum number of topics set to 36, and default values for other parameters.

### LITE and NITE modelling with MIRA

We employed the LITE and NITE models in MIRA ^51^ to characterize the regulatory influence of chromatin accessibility on gene expression. The local chromatin accessibility-influenced transcriptional expression (LITE) model focused on cis-regulatory elements (CREs) surrounding the gene transcriptional start site (TSS). For each gene locus, the regulatory effects of nearby accessible peaks were modeled to decay exponentially with distances from the TSS, with separate decay rates learned for upstream and downstream regions. Non-redundant human TSS annotations (hg38 GENCODE VM39) were used to assign unique TSS to each gene symbol. The LITE model was instantiated using the ‘LITE_Model’ function with default parameters for highly variable genes with annotated TSS (n=3253). Pretrained RNA and ATAC topic models were referenced. Raw expression and accessibility data were fitted to optimize expression estimates for each gene based on its local accessibility state in each cell. 3150 out of 3253 genes were fitted successfully.

The nonlocal chromatin accessibility-influenced transcriptional expression (NITE) model extends the LITE model by incorporating the genome-wide accessibility landscape captured by the ATAC topics. The ‘spawn_NITE_model’ function was used to initialize the NITE model with the same topic model references, genes, and pretrained parameter values from the LITE model. NITE models for 3120 out of 3150 genes were fitted successfully.

We next compared the predictive capacity between LITE and NITE models to determine the extent to which gene expression was driven by local chromatin accessibility. Cases where the NITE model provided a superior fit compared to the LITE model suggest that local chromatin features alone are not sufficient to predict gene expression, indicating a potential role for distal or global regulatory mechanisms. The ‘get_NITE_score_cells’ function was used to calculate cell-level NITE score, a one-number metric quantifying the statistical divergence between expression and local accessibility for each cell. A high NITE score indicates cell states where gene expression is less closely tied to local chromatin accessibility.

### Gene Regulatory Networks (GRNs)

To build gene regulatory networks, we combined different functions from Scanpy 113 and SCENIC+ ^80^.

#### GRNs pre-processing

Clusters and FA projection from multiVI were used as inputs for the preprocessing steps. Highly variable genes were selected using sc.pp.highly_variable_genes function. For the single-cell ATAC sequencing (scATAC-seq) data, fragment file paths were specified in a dictionary pointing to the atac_fragments_.tsv.gz files. Chromosome size information for the hg38 human genome assembly was downloaded from the UCSC Genome Browser using the requested library. This data was loaded into a pandas DataFrame with columns for chromosome (Chromosome), start position (Start), and end position (End). The chromosome size information was converted into a PyRanges object for genomic range operations. Pseudobulk profiles were generated using the pycisTopic library’s export_pseudobulk function. This involved aggregating scATAC-seq fragments by clusters, specified in the cell_data DataFrame. MACS2 was used for peak calling on pseudobulk BED files. Parameters were set for narrow peak calling, including the genome size (hs), shift size (73), extension size (146), and a q-value threshold of 0.05 for peak significance. Consensus peaks were identified across clusters by analyzing overlaps among narrow peaks. Transcription start site annotations were retrieved from the Ensembl database using Pybiomart, focusing on protein-coding transcripts. Quality control metrics were computed based on these annotations and the scATAC-seq fragment data. Metrics included unique fragment counts, duplication rates, insert size distributions, and fraction of reads in peaks (FRIP). Quality control thresholds were applied to filter for high-quality cells. A cistopic object, specific for integrative analysis of ATAC-seq data, was generated using the pycisTopic library. This object was created by specifying the paths to the ATAC-seq fragments, consensus regions, the blacklist file, cell barcode metadata, and the subset of valid barcodes passing the filters. The cistopic object was further enriched with cell metadata from the scRNA-seq analysis, allowing for integrative analyses of transcriptomic and epigenomic data. The object was serialized using the Python pickle module for persistence and subsequent analyses. Latent Dirichlet Allocation (LDA) models were fitted to the cistopic object to infer the underlying topics across a range of specified topic numbers (n_topics=[2,4,10,16,32,48]). The fitting process was configured with specific parameters for iteration count (n_iter=500), parallel processing (n_cpu=5), randomness control (random_state=555), and hyperparameters (alpha=50, eta=0.1). The models were saved for persistence and later loaded for evaluation. The best-fitting model was selected from the models with 16 topics for downstream analysis.

#### GRNs analysis

The python pickle module was used to retrieve serialized objects obtained from the previous preprocessing steps. Specifically, we worked with LDA models stored in “scATAC/models/Sample_ID_models_500_iter_LDA.pkl” and a cistopic object from “scATAC/cistopic_obj.pkl”. To binarize the topics, we applied Otsu’s method and selected the top 3000 topics using the “Top 3k” approach. For motif analysis, we utilized PyCistarget, leveraging specific databases and annotations. These included motif rankings (hg38_screen_v10_clust.regions_vs_motifs.rankings.feather), motif scores, (hg38_screen_v10_clust.regions_vs_motifs.scores.feather) and motif annotations (motifs-v10nr_clust-nr.hgnc-m0.001-o0.0.tbl), all available from Aerts lab resources. Within the SCENICPLUS framework, we integrated enhancer-to-gene relationships. Additionally, we examined transcription factor (TF) to gene relationships by referencing a curated list of transcription factors from the University of Toronto database (utoronto_human_tfs_v_1.01.txt). The GRN Construction process involved using the build_grn function from the scenicplus.grn_builder.gsea_approach module. The function was configured with key parameters such as a minimum of 10 target genes per GRN, an adjusted p-value threshold of 1, and a minimum of 0 regions per gene considered in the GRN. Quantiles for filtering were set to 0.85, 0.90, and 0.95, and the top numbers of regions to genes per gene and per region were set to (5, 10, 15) and (), respectively. The GRNs were binarized using BASC with dichotomization of TF-to-gene, region-to-gene, and eRegulon correlations set to true, and a rho threshold of 0.05. Extended motif annotations were retained, and eRegulons were merged based on their importance. The order of regions to genes and TFs to genes was based on their importance. The eRegulons_importance key was added for identification, with the cistromes key set to ‘Unfiltered’. To prepare the data for scoring based on eRegulon signatures, we employed the get_eRegulons_as_signatures function. Scoring was performed across chromatin and transcriptome layers using the make_rankings and score_eRegulons functions from the scenicplus.cistromes module. Our evaluation focused on eRegulon activity across different regions and genes. The parameters included the use of region and gene ranking for target specification, an AUC threshold of 0.05 for enrichment significance, and scoring was performed without normalization, utilizing plate_number_1 for computation. The TF_cistrome_correlation function analyzed the correlation between TFs and eRegulons based on the clusters variable, focusing on both gene-based and region-based signatures.

### Lentiviral shRNA preparation and cell transduction

Two gene-targeting shRNAs were selected and compared with a control non-targeting shRNA (Ctrl).

#### Lentivirus preparation

Lentiviral particles were produced by combining a packaging plasmid (psPAX2) and an envelope-coding plasmid (PMD2.G) with vectors containing the desired targeting DNA sequences ^46^. HEK 293 packaging cells were transfected using calcium phosphate reagent, at a ratio of 1:0.75:0.25 for transfer, packaging, and envelope plasmids, respectively. Viral particle supernatants were harvested at 48 and 72 h post-transfection and concentrated by ultracentrifugation at 50,000 x g for 2 hours at 4°C.

#### Lentiviral infections

CD34-positive cells were pre-stimulated for 24 hours in complete differentiation medium (DM), supplemented with cytokines. After 24 hours, cells were infected with lentiviral particles at an MOI of 50, in the presence of 4 µg/ml hexadimethrine bromide (Polybrene) and spinoculated at 800 x g for 30 minutes at RT to increase infection efficiency ^46^. Two rounds of lentiviral infection at 24-hour interval were performed. Cells were collected 48h post-transduction, and the efficiency of knockdowns and the expression of target genes were assessed by RT-qPCR analysis.

### Effector TF ChIPseq signal over eRegulon target CREs

RPKM-normalized ChIPseq coverage tracks for ERG and RUNX1 in HSC ^91^ were obtained from GEO: GSE231422. Merged coverage for immunoglobulin G (IgG) in HSC, CMP, GMP, and MEP was also downloaded as a control. Heatmap and composite profiles showing ERG and RUNX1 ChIPseq signal over predicted target CREs within the ERG and RUNX1 eRegulons were plotted respectively using deepTools v3.5.4.

### Overlap between genes essential for erythroid differentiation and erythroid-specific GRNs

277 genes required for terminal erythroid differentiation identified through genome-scale CRISPR knock-out screen (Log2 fold change < −1 and false discovery rate < 0.01 ^95^ were obtained from https://cdb-rshiny.med.umich.edu/Khoriaty_Erythropoiesis/. We filtered 21 microRNAs and 8 genes not detected in our TEAseq data, leaving 248 genes considered in downstream analyses. These 248 genes essential for erythroid differentiation were overlapped with our erythroid-specific GRNs, including effector TFs and their predicted target genes within BFU-e, CFU-e1, CFU-e2, ProEB, BasoEB and Poly-OrthoEB eRegulons.

